# Orc4 spatiotemporally stabilizes centromeric chromatin

**DOI:** 10.1101/465880

**Authors:** Lakshmi Sreekumar, Kiran Kumari, Asif Bakshi, Neha Varshney, Bhagya C. Thimmappa, Krishnendu Guin, Leelavati Narlikar, Ranjith Padinhateeri, Rahul Siddharthan, Kaustuv Sanyal

## Abstract

Spatiotemporal regulation in DNA replication maintains kinetochore stability. The epigenetically regulated centromeres (CENs) in the budding yeast *Candida albicans* have unique DNA sequences, replicate early and are clustered throughout the cell cycle. In this study, the genome-wide occupancy of replication initiation protein Orc4 reveals its abundance at all CENs in *C. albicans*. Orc4 associates with four different DNA motifs, one of which coincides with tRNA genes. Hi-C combined with genome-wide replication timing analyses identify enriched interactions among early or late replicating Orc4-bound regions. A simulated polymer model of chromosomes reveals that early replicating and strongly enriched Orc4-bound sites localize towards the kinetochores. Orc4 is constitutively localized to CENs, and both Orc4 and Mcm2 stabilize CENPA. CENPA chaperone Scm3 localizes at the kinetochore during anaphase, coinciding with the loading time of CENPA. We propose that this spatiotemporal nuclear localization of Orc4, with Mcm2 and Scm3, recruits CENPA and stabilizes centromeric chromatin.

## Introduction

The timely duplication of the genetic material and faithful chromosome segregation maintains genome stability. In eukaryotes, the origin recognition complex (ORC) comprising of Orc1-6 along with Cdc6, Cdt1 and the minichromosome maintenance complex (Mcm2-7) initiate DNA replication at multiple discrete sites on a chromosome that serve as DNA replication origins (Leonard and Mechali 2013). ORC recognizes its cognate binding sites in a species-specific manner – an AT-rich consensus sequence in the budding yeast *Saccharomyces cerevisiae* (Wyrick, Aparicio et al. 2001) to AT-rich asymmetric sequences in the fission yeast *Schizosaccharomyces pombe* (Chuang and Kelly 1999) while its binding is non-specific to any DNA sequence in humans (Vashee, Cvetic et al. 2003). The robust programing of the time of firing of replication origins during S phase is required to ensure the availability of the limiting concentration of replication initiation factors (Aparicio 2013). Genomic regions that replicate early during S phase have a higher propensity to attract ORC and are more efficient in origin firing, whereas late replicating regions appear to be more stochastic contributing to inefficient initiation events (Mesner, Valsakumar et al. 2011). The spatial organization of chromosomes within the nucleus also favors the accessibility of initiation factors to complete DNA replication in a timely manner (Aparicio 2013). Chromosome conformation capture (3C) analysis has revealed higher interaction frequencies between early origins in *S. cerevisiae* (Duan, Andronescu et al. 2010). Evidently, understanding the topology of the three-dimensional (3D) genome and the factors that spatiotemporally regulate the chromosomal architecture is necessary to explain how the genome is organized into various functional domains.

The faithful segregation of the duplicated genome is facilitated by centromeres (CENs) which act as the binding platform for kinetochore proteins. In most eukaryotes, CENs are specified by a CEN-specific histone H3 variant, CENPA through epigenetic mechanisms, where the underlying DNA sequence appears dispensable for CEN establishment (Malik and Henikoff 2009). The epigenetic specification of CENs in most eukaryotes is evident as CEN DNA are the most rapidly evolving loci of the genome (Malik and Henikoff 2009). The targeted loading of CENPA is temporally separated from bulk H3 chromatin assembly in the cell cycle and is mediated by the specific chaperone Holliday junction recognition protein (HJURP) (Kato, Sato et al. 2007) and its analogs in various species. The yeast homolog Scm3 loads CENPA during S phase in *S. cerevisiae* (Williams, Hayashi et al. 2009), whereas in *S. pombe* CENPA loading occurs during G2 (Shukla, Tong et al. 2018). The functional homolog of HJURP in *Drosophila*, CAL1 loads CENPA during late telophase (Dunleavy, Beier et al. 2012). In humans, the G1 loading of CENPA by HJURP (Foltz, Jansen et al. 2009) is regulated by the Mis18 complex (Barnhart-Dailey, Trivedi et al. 2017). In *S. pombe*, RNAi and heterochromatin are required for CEN establishment (Folco, Pidoux et al. 2008). Unlike metazoa, most fungi have early replicating CENs (Raghuraman, Winzeler et al. 2001, Kim, Dubey et al. 2003, Koren, Tsai et al. 2010). Vertebrate CENs replicate between mid-late S phase (Ten Hagen, Gilbert et al. 1990). CENPA nucleosome disruption following DNA replication transiently creates gaps or nucleosome-free regions which have to be reassembled for CEN propagation. The placeholder model proposes that the gaps generated upon parental CENPA eviction are occupied by placeholder molecules like H3 in *S. pombe* (Shukla, Tong et al. 2018) and H3.3 in *Drosophila* (Dunleavy, Almouzni et al. 2011). Hence, CENPA replenishment by its replication-independent loading is necessary for its maintenance.

Once established, CENPA chromatin can epigenetically self-propagate through multiple cell divisions, best studied in the case of ectopic or neocentromere activation at non-centromeric loci upon native CEN inactivation (Warburton 2004). However, mechanisms contributing to maintenance of centromeric chromatin is relatively unclear. Growing evidence suggests a role of replication initiation proteins in CEN function (Natsume, Muller et al. 2013). Fungal CENs replicate early in S phase (Pohl, Brewer et al. 2012) to maintain kinetochore integrity and pericentromeric cohesion (Kitamura, Tanaka et al. 2007, Natsume, Muller et al. 2013). In humans, Orc2 localizes to CENs (Prasanth, Prasanth et al. 2004), and HJURP along with Mcm2 is required for CENPA inheritance during S phase (Zasadzinska, Huang et al. 2018). *In vitro* experiments suggest that Mcm2 and Asf1 cochaperone dimers of H4 with both the canonical and variant forms of H3 through its histone-binding mode (Huang, Stromme et al. 2015, Richet, Liu et al. 2015). Recently, DNA replication has been implicated to remove ectopically loaded non-centromeric CENPA for the precise reloading of centromeric CENPA during G1 in humans (Nechemia-Arbely, Miga et al. 2019). Also, replication fork termination seen at CEN promotes CEN DNA loop formation which is ultimately required for kinetochore assembly (Cook, Bennett et al. 2018). Hence, there is an implicit crosstalk of replication initiation proteins for CEN function.

In yeast, CENs of all chromosomes cluster close to the spindle pole body (SPB) embedded into the nuclear membrane and establish inter-chromosomal interactions as shown by 3C experiments (Jin, Fuchs et al. 2000, Duan, Andronescu et al. 2010). Additionally, computational models for the *S. cerevisiae* and *S. pombe* genomes based on highly structured Hi-C contact maps have revealed strong CEN clustering and significant but weaker telomere (TEL) interactions along the nuclear envelope (Tjong, Gong et al. 2012, Gong, Tjong et al. 2015). The clustered CENs are present in a compact chromatin environment both in vertebrates (Nishimura, Komiya et al. 2018) and in the budding yeast *Candida albicans* (Sreekumar, Jaitly et al. 2019). *C. albicans* is one such organism where CENPA recruitment is not defined by a consensus DNA sequence, but instead by an early replicating 3-5 kb unique domain on every chromosome present in a cluster (Sanyal, Baum et al. 2004, Baum, Sanyal et al. 2006, Koren, Tsai et al. 2010). Centromeric chromatin is more compact and exhibits strong *trans* inter-centromeric interactions as shown by genetic experiments and Hi-C analyses in *C. albicans* (Sreekumar, Jaitly et al. 2019). It can efficiently activate neocentromeres at CEN-proximal regions upon the deletion of a native CEN (Thakur and Sanyal 2013). Deletion of CEN-proximal replication origins abrogates CEN function and debilitates kinetochore stability in this organism (Mitra, Gomez-Raja et al. 2014). Replication fork stalling at CENs is facilitated by the homologous recombination (HR) proteins, Rad51 and Rad52, which are known to physically interact with CENPA, stabilizing the kinetochore (Mitra, Gomez-Raja et al. 2014). Having a constitutive kinetochore ensemble (Thakur and Sanyal 2012), new CENPA in *C. albicans* is known to load during anaphase (Shivaraju, Unruh et al. 2012). Surprisingly, depletion of an essential kinetochore protein, irrespective of its location at the kinetochore, disintegrates the kinetochore ensemble leading to degradation of CENPA by ubiquitin-mediated proteolysis (Thakur and Sanyal 2012). The factors that stabilize CEN chromatin in *C. albicans* in the absence of complete RNAi machinery, typical pericentric heterochromatin and factors such as the Mis18 complex remain an enigma.

In the present study, we report factors that help maintain and regulate the short epigenetically regulated centromeric chromatin in *C. albicans*. Our genome-wide binding analysis of Orc4 reveals a constitutive localization of Orc4 at all CENs, which is pertinent for CENPA stability. Categorizing the Orc4-bound sites according to replication time zones led to a strong correlation between the replication timing and spatial interaction frequencies of the Orc4-bound regions in the genome. The computational polymer model of chromosomes developed in this study, the first to be reported for *C. albicans*, demonstrates the spatial distribution of Orc4 within the nucleus based on replication timing. We also observe that CENPA is loaded to CEN by the chaperone Scm3 during anaphase and is stabilized by Orc4 along with Mcm2, leading us to finally propose a model for establishment and maintenance of centromeric chromatin in *C. albicans*.

## Results

### Orc4 binds to discrete regions uniformly across the *C. albicans* genome

To examine the replication landscape of the *C. albicans* genome, we sought to determine the genome-wide occupancy of Orc4. Orc4 in *C. albicans* is a 564-aa long protein (Padmanabhan, Sanyal et al. 2018) that contains the evolutionarily conserved AAA+ domain (Walker, Saraste et al. 1982) (Figure S1A). We raised polyclonal antibodies in rabbits against a peptide from the N-terminus of the native Orc4 (aa 20-33) of *C. albicans* (Figure S1B). Western blot of the whole cell extract of *C. albicans* SC5314 *(ORC4/ORC4*) yielded a strong specific band at the expected molecular weight of approximately 64 kDa when probed with purified anti-Orc4 antibodies (Figure S1C). Indirect immuno-fluorescence microscopy using anti-Orc4 antibodies revealed that Orc4 was strictly nuclear localized at all stages of the *C. albicans* cell cycle (Figure 1A), a feature of the ORC proteins conserved in *S. cerevisiae* as well (Dutta and Bell 1997).

**Figure 1.**
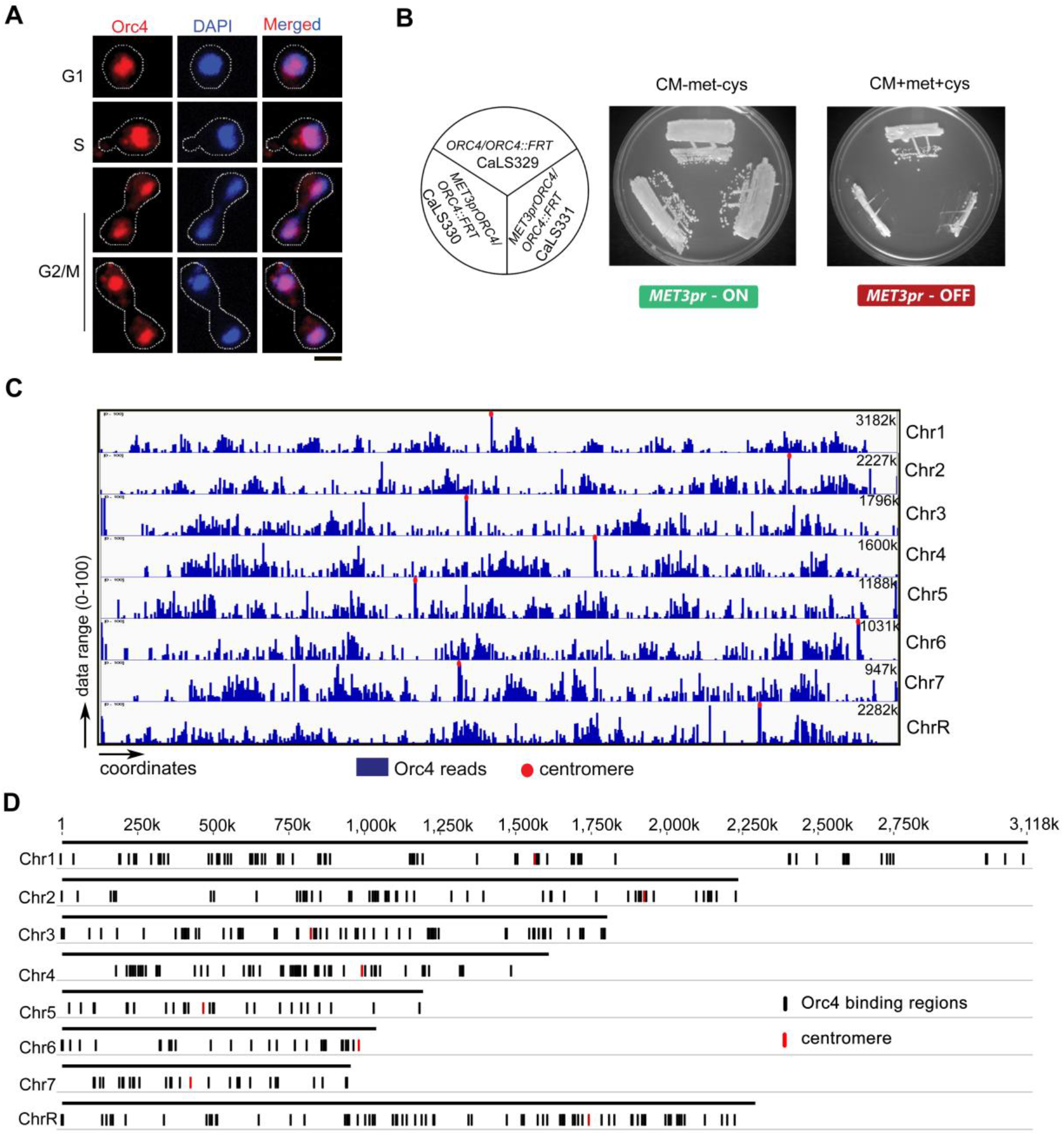
Orc4, an essential subunit of the origin recognition complex, is nuclear localized and binds to discrete loci in the *C. albicans* genome. (**A**) Nuclear localization of Orc4 in *C. albicans* SC5314 cells as evidenced by staining with anti-Orc4 antibodies (red) and DAPI (blue). Bar, 5 μm. (**B**) The promoter of *MET3* in *C. albicans*, expressed in the absence of methionine and cysteine and repressed in the presence of both, was used for the conditional expression of *ORC4*. CaLS329 (*ORC4*/*ORC4*::*FRT)* with one deleted copy of *ORC4*, and two independent transformants, CaLS330 and CaLS331 (*MET3prORC4/ORC4*::*FRT*), where the remaining wild-type copy was placed under the control of the *MET3* promoter were streaked on plates containing inducible (CM-met-cys) or repressible (CM+ 5 mM cys + 5 mM met) media and photographed after 48 h of incubation at 30°C. (**C**) ChIP-sequencing analysis revealed that Orc4 binds to discrete genomic sites in *C. albicans*. The total Orc4 reads (blue histogram) were obtained by subtracting the relative number of sequencing reads from the whole cell lysate from the Orc4 ChIP sequence reads and aligning them to the reference genome *C. albicans* SC5314 Assembly 21. Red dots indicate CENs. (**D**) Orc4 binding regions (black) on each of the eight *C. albicans* chromosomes including all eight CENs (red).

Orc4 is an evolutionarily conserved essential subunit of ORC across eukaryotes (Chuang and Kelly 1999, Dai, Chuang et al. 2005). A conditional mutant of *orc4* in *C. albicans* constructed by deleting one allele and replacing the endogenous promoter of the remaining *ORC4* allele with the repressive *MET3* promoter of *C. albicans* (Care, Trevethick et al. 1999), was unable to grow in non-permissive conditions (Figure 1B). Hence, Orc4 is essential for viability in *C. albicans.* We confirmed the depletion of Orc4 protein levels from the cellular pool by performing a western blot analysis in the Orc4 repressed as compared to expressed conditions (Figure S1D). Subsequently, we used the purified anti-Orc4 antibodies as a tool to map its binding sites across the *C. albicans* genome. Orc4 ChIP sequencing in asynchronously grown cells of *C. albicans* yielded a total of 417 discrete Orc4 binding sites with 414 of these belonging to various genomic loci while the remaining three mapped to mitochondrial DNA (Figures 1C, 1D). We validated one region on each of the eight chromosomes by Orc4 ChIP-qPCR (Figure S1E). Strikingly, all CENs were found to be strongly enriched with Orc4. While the majority of the binding loci (>300) spanned ~1 kb in length, all eight CENs had an Orc4 occupancy spanning 3-4 kb (Figure S1F). Approximately 61% of the Orc4 binding regions in our study were present in the gene bodies (252/417) of *C. albicans*.

### Orc4 displays differential DNA binding modes which are spatiotemporally positioned in the genome

We used the *de novo* motif discovery tool DIVERSITY (Mitra, Biswas et al. 2018) on the *C. albicans* Orc4 binding regions. DIVERSITY allows for the fact that the profiled protein may have multiple modes of DNA binding. Here, DIVERSITY reported four binding modes (Figure 2A). The first mode, mode A is a strong motif GAnTCGAAC, present in 50 such regions, 49 of which were found to be located within tRNA gene bodies and one within the tDNA regulatory region. The other three modes were low complexity motifs, TGATGA (mode B), CAnCAnCAn (mode C) and AGnAG (mode D). Strikingly, each of the 417 binding regions was associated with one of these motifs. Mode C has been identified in a previous study (Tsai, Baller et al. 2014) in which ORC binding sites in the *C. albicans* genome were mapped using a microarray-based approach. ORC binding regions of these two studies share the maximum overlap at the mode A ORC containing sites (Figure S2A). Taken together, these results suggest that Orc4 in *C. albicans* is not specified by a single DNA binding site, rather displays differential DNA binding modes.

**Figure 2.**
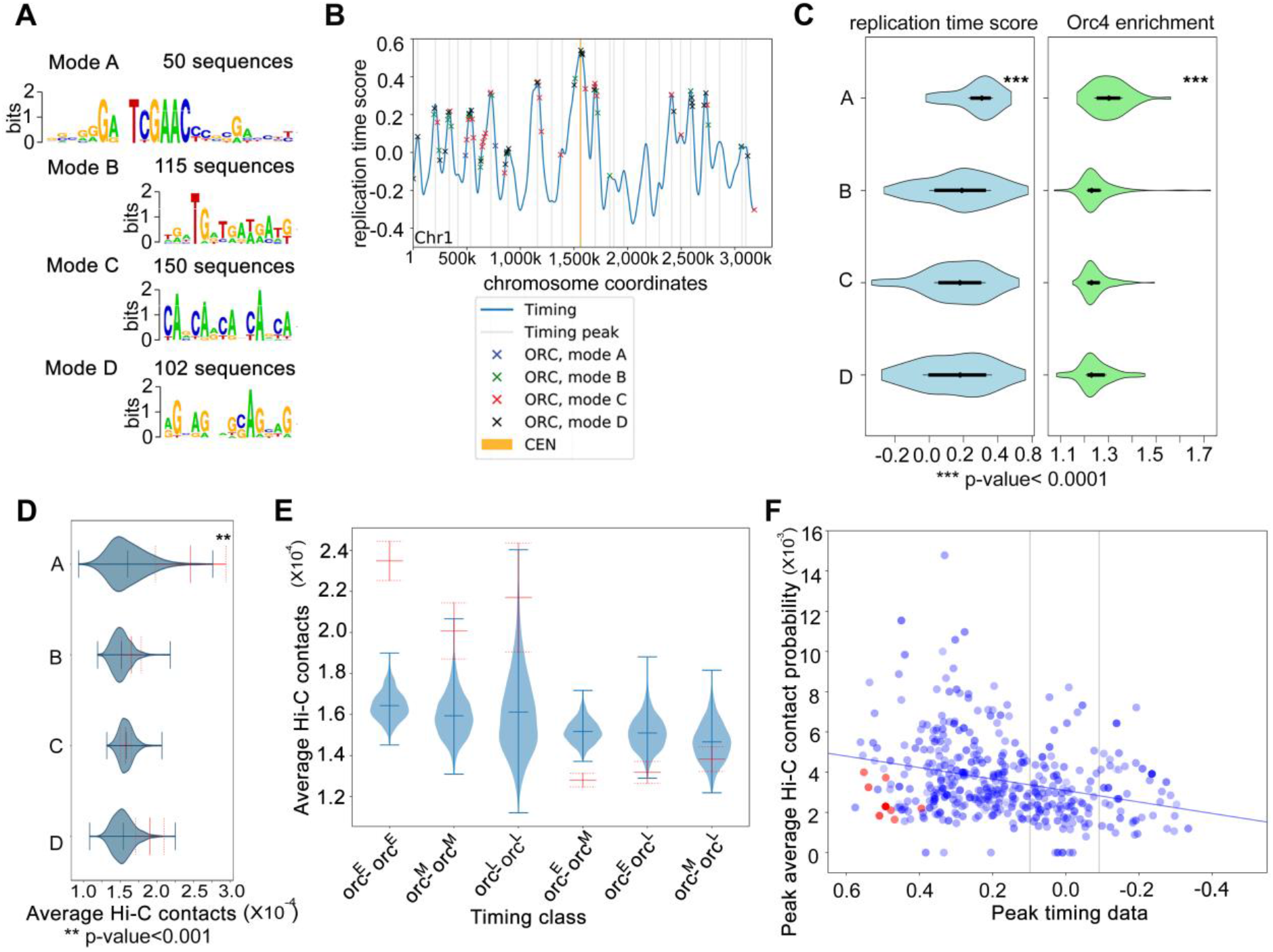
Orc4 is associated to four different DNA binding modes which are spatiotemporally positioned across the genome. (**A**) The four different modes identified by DIVERSITY (A, B, C, D) and their distribution across the 417 Orc4 binding regions have been listed. (**B**) Orc4 ChIP-seq peaks denoted as asterisks, colored according to the four modes identified by DIVERSITY, are overlaid on the replication timing profile of Chr1 in *C. albicans* from a previous study (Koren, Tsai et al. 2010). Higher time score indicates an early time of replication. Color-coded stars indicate each of the four motifs identified by DIVERSITY which covers all the 414 chromosomal sites. Light grey lines indicate local maxima in replication time. CENs are shown in yellow. (**C**) Violin plots depicting the replication timing scores (Koren, Tsai et al. 2010) (blue) and Orc4 enrichment (green) of all the Orc4 peaks classified according to each of the four modes. **(D**) Average pairwise interaction scores (Burrack, Hutton et al. 2016) of Orc4 peaks with other peaks in the same mode. Solid red = mean, dotted red = standard error, violins are from 1,000 sets of randomised data (randomly selected genomics regions with the same size and chromosomal distribution as the peaks in that mode). **(E)** Mean Hi-C interactions (solid red) with standard error (dotted red) within and across each of the three timing classes (orc^E^, orc^M^ and orc^L^). These indicate higher interaction values within orc^E^ and within orc^L^ domains. Blue violins indicate mean interactions across 1,000 randomizations, as in (D). (**F**) A scatter plot of Hi-C contacts, replication timing and fold enrichment values of Orc4 binding regions. Each dot is an individual Orc4 peak, with its color intensity corresponding to its ChIP-seq enrichment value; red dots are peaks overlapping the eight CENs. The *y*-axis (peak average Hi-C contacts) represents the average of 10 best contacts for each peak of Orc4.

To categorize the replication timing of Orc4 binding sites, we utilized the available fully processed replication timing profile of the *C. albicans* genome (Koren, Tsai et al. 2010). Sorting the replication data based on replication time of the entire genome, the first one-third (33.3%) of the replicating regions was classified as early, the second one-third regions were classified as mid and the remaining were late replicating regions. Comparing the Orc4 sites to this profile, we found 218 early or orc^E^ sites (~52% of the total), 127 mid or orc^M^ sites (~30%) and 69 late or orc^L^ sites (~16%). We then overlaid the DIVERSITY motifs onto the timing profile (Figures 2B, S2B). We observed a significant advanced replication timing of the tRNA associated motifs (mode A) (Figure 2C). The other three modes (B, C, D) display no significant bias towards an early replication score. Moreover, we could correlate early replication timing with an increased enrichment of Orc4 in these regions (Figure 2C). In addition, all the motifs were located towards the local maxima of the timing peaks.

To locate these regions within the nuclear territory, we mapped the interactions made by the Orc4 binding regions with each other using the Hi-C data from a previous study in *C. albicans* (Burrack, Hutton et al. 2016). All the Orc4 binding regions were aligned in an increasing order of their replication timing (early to late) and subsequent interactions were mapped. Similar analysis was performed for the whole genome of *C. albicans.* We observed that the overall “only-ORC” interactions were higher than the whole-genome “all” interactions, suggesting that Orc4 binding regions interacted more than the genome average (Figures S2C, S2D). Hi-C analysis also revealed that the mode A sites formed stronger interactions among themselves than all the other modes (Figure 2D). We also performed a comparative analysis of the contact probabilities of the mode A sites with the rest of the tRNA genes in the genome, and observed a significantly higher interaction of mode A tRNAs over the rest (Figures S2E, S2F). Additionally, there was a significant increase in interaction frequencies between similarly timed domains (orc^E^ – orc^E^; orc^M^ – orc^M^; orc^L^ – orc^L^) as compared to interactions across domains (Figure 2E). Upon arraying the Orc4 peaks according to their replication timing scores reported previously (Koren, Tsai et al. 2010) against the average Hi-C interaction frequencies, we could observe a weak but significant correlation between contact probability and replication timing (Figure 2F). We also found higher Orc4 enrichment in the orc^E^ – orc^M^ regions as the majority of the Orc4 peaks were located in the middle of the pack (Figure 2F). Taken together, our analyses suggest that Orc4-bound regions with a similar timing in replication tend to associate together. In addition, replicating time zones may be specified by the availability of the replication initiation factors at various nuclear territories.

### Polymer modeling of *C. albicans* chromosomes reveals replication timing driven positioning of Orc4 within the nucleus

Hi-C analysis alone does not reveal the positioning of a particular locus within the nucleus. These intra- and inter-chromosomal interaction frequencies can be converted to linear distance approximations and averaged across populations to generate computational models that yield an ensemble of genomic conformations (Berger, Cabal et al. 2008, Gursoy, Xu et al. 2017). To study the 3D structural organization of the *C. albicans* genome, we resorted to polymer modeling of chromosomes using the contact probability data from the published Hi-C experiment (Burrack, Hutton et al. 2016). To do this, we used a statistical approach where each bead-pair is either bonded or not bonded based on the available contact probability data (Table S1). At first, we compared the contact probability data obtained from our simulation of 1,000 different configurations (Figure S3A) with the Hi-C experimental data with a resolution of 10 kb (Figure S3B) to ensure that our simulation had indeed satisfactorily recovered the input contact matrix. Contact probability for a bead-pair (i,j) from the simulation is calculated by averaging the bonding function bij over 1,000 realizations. The function b_ij_=1, in the case of a contact (r_ij_<1.5 l_0_) and zero otherwise. Here, the r_ij_ is the spatial distance between bead i and bead j and l_0_ is the natural extension of the connector spring. The contact probability from the simulation was found to be in close agreement with the Hi-C data showing the reliability of the model. From our simulations, we could predict the average spatial distance between any two beads within the genome. For a given contact probability, the corresponding average spatial distance could be computed (Figure S3C). We fixed the position of one of the CEN beads as the reference, and hence sought to determine the 3D location of each of these beads.

To examine the genome-wide distribution of Orc4 in the 3D nuclear space, we mapped the Orc4 ChIP-seq data to the corresponding coarse-grained beads. Using our simulation, we computed 3D locations of the Orc4 binding sites located in the early, mid and late replicating regions of the genome (as categorized previously). From the experimentally obtained contact probability, it was observed that orc^E^ sites interact strongly with CENs as compared to orc^M^ and orc^L^ sites (Figure 3A). Our simulations show the corresponding distances between the above-mentioned regions, where the average distance between the orc^E^ sites with CENs is significantly shorter than the average distance between CEN and orc^M^/orc^L^ regions (Figure 3B). To visualize the location of binding sites of orc^E^, orc^M^ and orc^L^ with respect to CENs and TELs, we chose one random configuration from an ensemble of 1,000 configurations. The orc^E^ regions are relatively closer to CENs (Figure 3C) while the orc^M^ sites are farther away (Figure 3D) and orc^L^ are the farthest from CENs (Figure 3E). Hence, there is a replication time-driven spatial distribution of Orc4 along the chromosomes with the highest concentration near CENs that decreases towards TELs (Figure 3F). Taken together, our computationa model revealed that Orc4 is not randomly distributed in the nucleus of *C. albicans*, instead is largely suggestive of a specific spatial organization driven by the replication time of Orc4 occupied loci (Figure 3G, Video S1).

**Figure 3.**
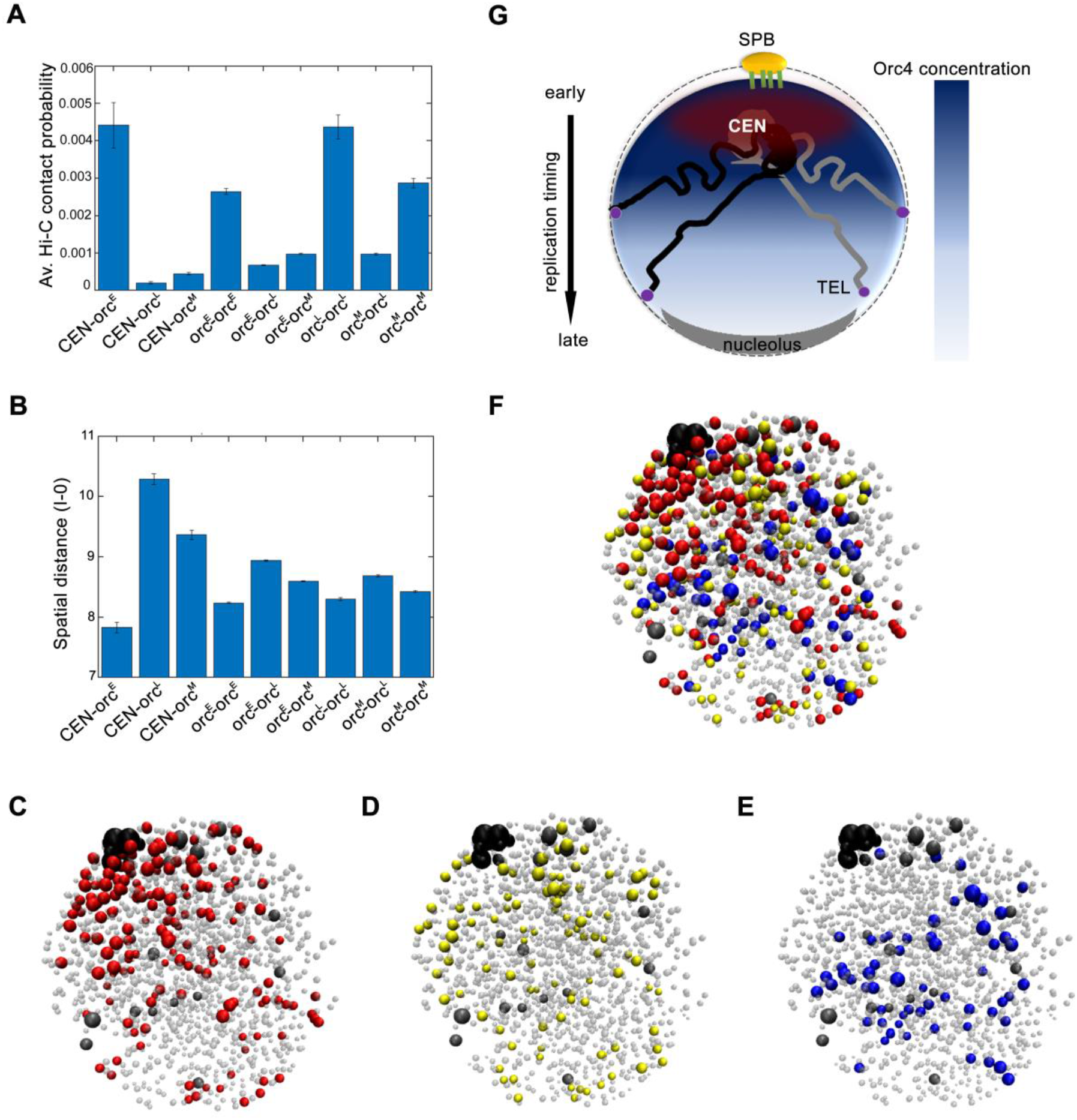
Early replicating Orc4 bound regions cluster around the clustered CENs. (**A**) The average contact probability for the indicated region for each of the timing class (across and within orc^E^, orc^M^ and orc^L^ domains) shows stronger CEN-orc^E^ interactions. (**B**) The average spatial distance between the indicated regions calculated from the Langevin simulation for each of the timing class (across and within the orc^E^, orc^M^ and orc^L^ domains) indicates shorter distances between CEN-orc^E^. (**C-F**) Snapshot of the 3D configuration of the *C. albicans* genome (1 out of 1,000 realizations) from the simulations shows all chromosomes in light grey, CENs in black and TELs in dark grey. Orc^E^ regions are shown in red (C), orc^M^ in yellow (D) orc^L^ in blue (E) and (F) represents all the Orc4 binding sites (orc^E^, orc^M^ and orc^L^). (**G**) Schematic of a budding yeast nucleus exhibiting a typical Rabl configuration where the clustered CENs are anchored near SPBs and TELs are often away from the CEN cluster and occasionally interacting with nuclear envelope. In *C. albicans*, the highest spatial enrichment of Orc4 is near the CEN cluster. Orc4 concentration gradually diminishes towards the opposite pole. Concomitantly, early replicating regions are located towards CENs and the late regions are towards TELs.

### Constitutive localization of Orc4 at CEN stabilizes CENPA

Since ORC is not known to be associated with CEN function, the strong enrichment of Orc4 at all CENs prompted us to examine its biological significance in *C. albicans*. Upon comparison of the Orc4 enrichment with CENPA occupancy in *C. albicans*, there was a striking overlap in the binding regions of these two proteins, indicating a strong physical association of Orc4 at CENs (Figures 4A, S4A, Table S2). To validate its role in CEN establishment, we examined its binding to *de novo* CEN formation at a non-native locus. *C. albicans* has been shown to efficiently activate neocentromeres at CEN-proximal regions when a native CEN is deleted (Thakur and Sanyal 2013). Orc4 was found to be enriched at the neocentromere hotspot *nCEN7-II* when the 4.5 kb CENPA-rich region on *CEN7* was deleted. Orc4 was not enriched at *nCEN7-II* in the wild-type strain with unaltered *CEN7* (Figure S4B). This result confirms that Orc4 helps in CEN establishment and further hints towards the possible role of replication initiator proteins in specifying CEN function in *C. albicans.*

**Figure 4.**
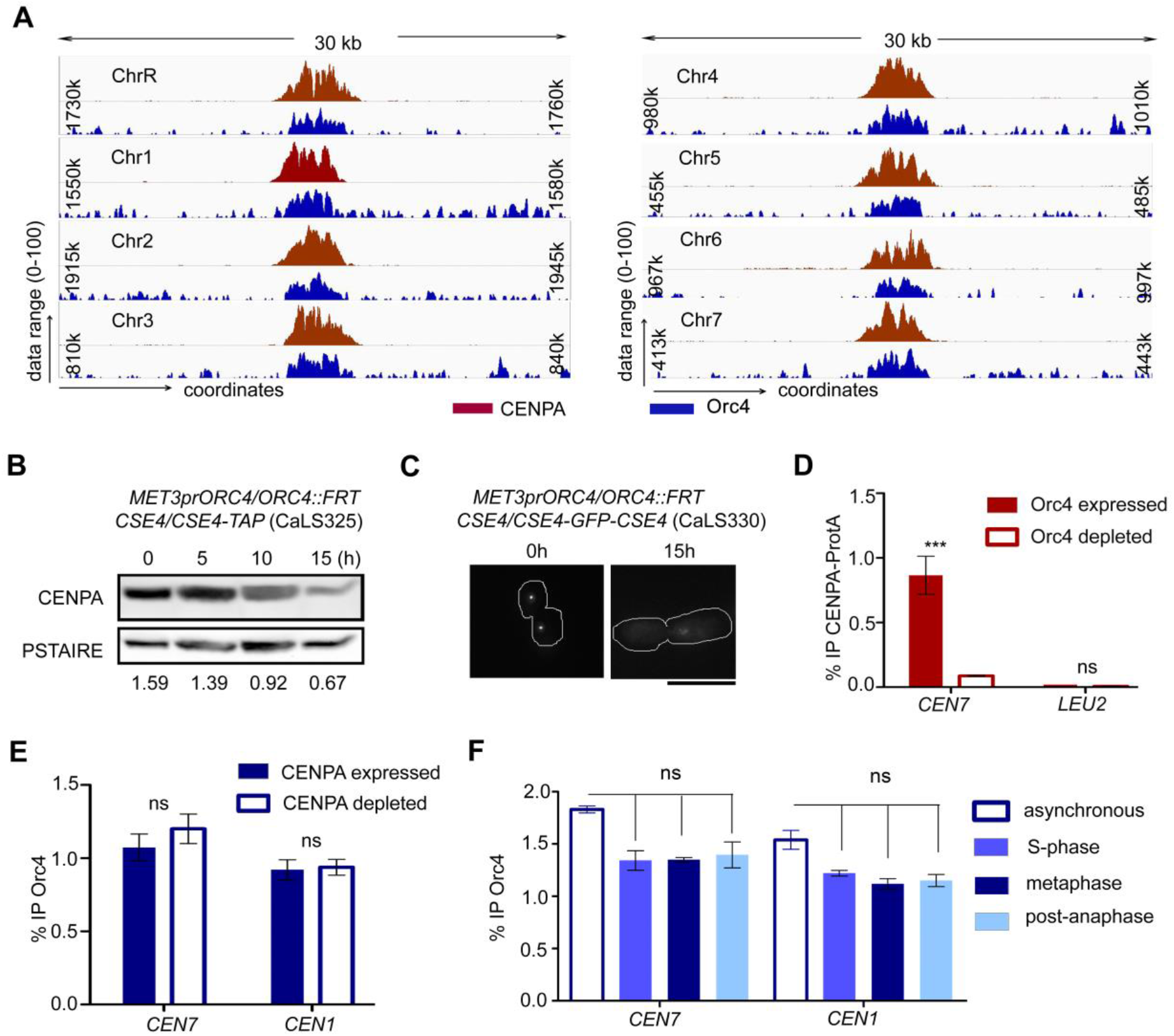
Centromeric localization of Orc4 stabilizes CENPA. (A) A 30 kb region harboring each CEN (*x*-axis) was plotted against the subtracted ChIP sequencing reads (*y*-axis) for CENPA (red) and Orc4 (blue). (**B**) Western blot of the whole cell lysate of CaLS325 (*METprORC4/ORC4::FRT CSE4/CSE4-TAP*) using anti-Protein A antibodies shows time-dependant decrease in CENPA levels upon Orc4 depletion when normalized with the loading control, PSTAIRE. (**C**) CENPA cluster delocalized upon a time course depletion of Orc4 in CaLS330. Scale bar, 10 μm. (**D**) ChIP-qPCR using anti-GFP antibodies revealed reduced CENPA enrichment at CEN upon Orc4 depletion in CaLS330 (*MET3prORC4/ORC4::FRT*) grown either in CM-met-cys or CM+5mM met + 5mM cys for 15 h. (**E**) Orc4 ChIP-qPCR revealed no significant reduction in centromeric Orc4 when CENPA was depleted for 8 h in YP with dextrose in the strain CAKS3b (*cse4/PCK1pr-CSE4*) (Sanyal and Carbon 2002). Two-way ANOVA was used to determine statistical significance. (**F**) ChIP-qPCR to show Orc4 enrichment in various stages of cell cycle: hydroxyurea treated (S phase), nocodazole treated (metaphase), *cdc15* mutant (Bates 2018) (post-anaphase). Percent IP values for Orc4 ChIP at CEN were normalized with non-centromeric regions enriched with Orc4. Each of the values were compared with an asynchronous culture control to determine statistical analysis using one-way ANOVA (***p<0.001, ns p>0.05).

To examine its role in CEN function, we assayed for CENPA localization in an *orc4* conditional mutant. Western blot analysis revealed a significant reduction of CENPA (Figure 4B) and declustering of the kinetochore (Figure 4C) upon prolonged Orc4 depletion. ChIP experiments revealed a 90% reduction in CENPA at CENs upon Orc4 depletion for 15 h reminiscent of kinetochore disintegration (Figure 4D). However, depletion of CENPA did not alter the levels of centromeric Orc4 (Figure 4E), indicating that Orc4 regulates CENPA localization at CENs but not vice-versa. We were intrigued to examine the centromeric occupancy of Orc4 at various stages of the *C. albicans* cell cycle. We determined the enrichment levels of Orc4 at CENs in S phase, metaphase and post-anaphase stages in comparison with an asynchronous culture to observe no significant difference across all stages (Figure 4F). These results suggested that Orc4 is constitutively localized at the kinetochore throughout the cell cycle in *C. albicans*. Spot dilution assays to determine the viability of *orc4* mutant after prolonged depletion revealed no observable drop in the viability of *orc4* mutant when grown in permissive media (Figure S4C) but an unsegregated kinetochore in more than 90% of the cells (Figure S4D). To rule out the possibility that a general replication stress can lead to loss of CENPA, we treated the cells with an S phase inhibitor, hydroxyurea (HU) and quantified the mean CENPA-GFP intensity in the strain YJB8675 (*CSE4-GFP-CSE4/CSE4)* (Joglekar, Bouck et al. 2008). Compared to an untreated control, we could not detect any significant difference in the CENPA-GFP intensity upon HU treatment (Figure S4E). This was further corroborated by performing a western blot analysis (Figure S4F) and ChIP-qPCR analysis (Figure S4G). These results together suggest that a global replication stress does not alter CENPA levels at CEN and constitutive Orc4 localization at CENs is essential for CENPA stability.

### Mcm2 affects CENPA stability and chromosome segregation

Apart from its canonical helicase activity during replication initiation, Mcm2 is known to bind to CENPA *in vitro* (Huang, Stromme et al. 2015). In order to examine its role in CENPA stability, we sought to characterize its homolog in *C. albicans. MCM2* is annotated as an uncharacterized ORF (orf19.4354) in the *Candida Genome Database* (candidagenome.org). BLAST analysis using *S. cerevisiae* Mcm2 as the query sequence revealed that orf19.4354 translates to a 101.2 kDa protein that contains the conserved Walker A, Walker B and the R finger motif together constituting the MCM box (Forsburg 2004) (Figures S5A, S5B). We tagged Mcm2 with Protein A at the C-terminus in BWP17 and then deleted the untagged allele of *MCM2* to generate a singly Protein A tagged strain CaLS334. Western blot analysis with the tagged protein lysate yielded a specific band at the expected molecular weight of 135 kDa, which could not be detected in the untagged lysate (Figure S5C). By indirect immuno-fluorescence microscopy, Mcm2-ProtA was found to colocalize with the nucleus in the G1 and S phases of the cell cycle (Figure 5A). However, in the large budded G2/M cells two distinct patterns of localization were observed. In the large budded cells with a single nucleus at the pre-anaphase stage, Mcm2 signal was weak or absent from the nucleus. We could localize Mcm2 in the nuclei of large budded cells where nuclear separation has occurred (post-anaphase) between the mother and daughter buds (Figure 5A), reminiscent of MCM proteins in *S. cerevisiae* (Yan, Merchant et al. 1993). We performed Mcm2 ChIP-qPCR analysis with primers corresponding to few of the Orc4 binding regions on each of the eight chromosomes and could detect four out of eight sites to be significantly enriched with Mcm2 over the control (*LEU2*) region (Figure S5D). Hence, a fraction of the Orc4 bound regions act as binding sites for Mcm2, hinting that these overlapping binding regions could be replication origins in *C. albicans*.

**Figure 5.**
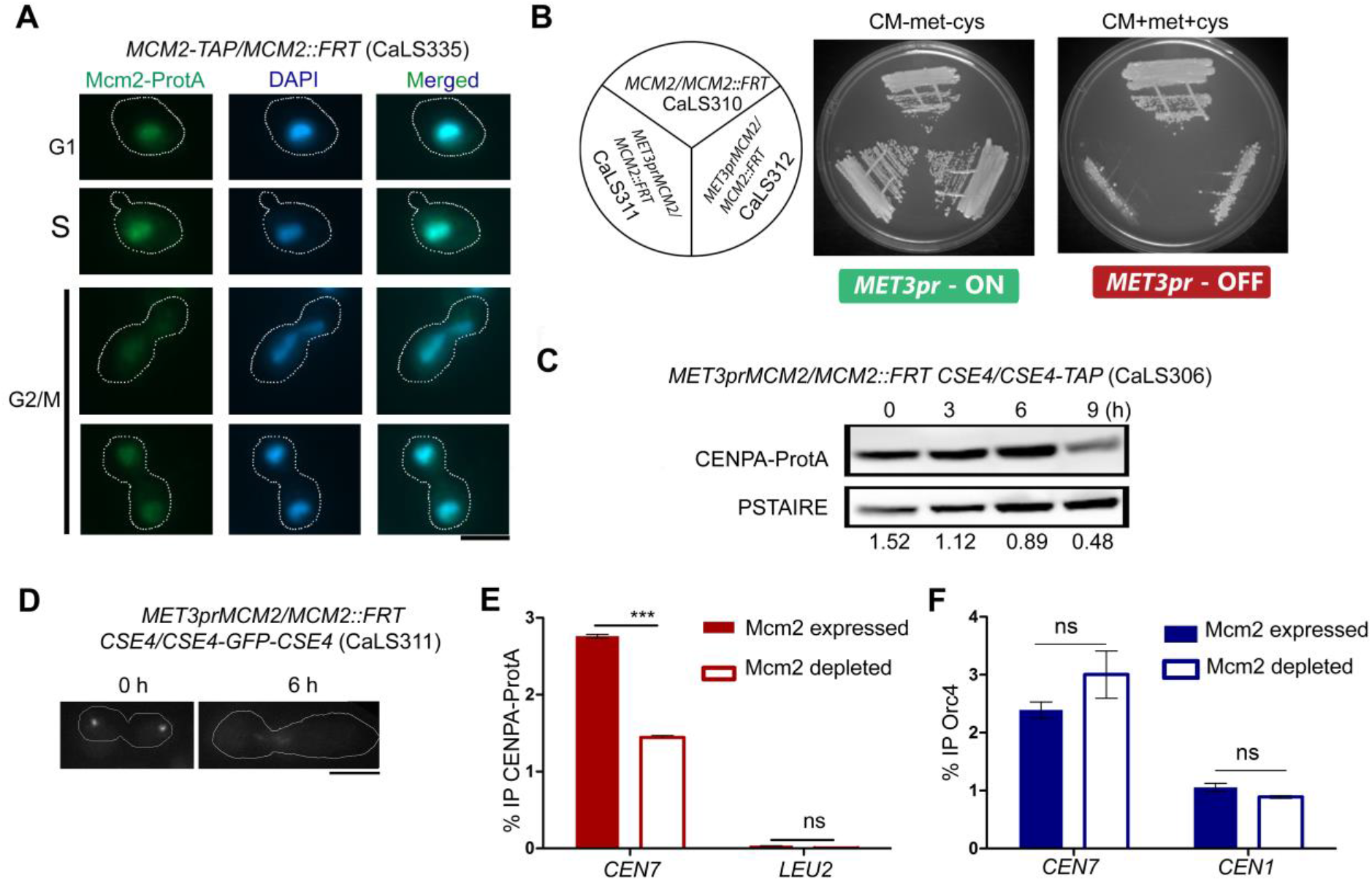
Mcm2 is essential for cell viability and CENPA stability in *C. albicans*. **(A)** Intracellular localization of Mcm2-ProtA in CaLS335 (*MCM2-TAP/MCM2::FRT*) cells stained with anti-Protein A antibodies (green) and DAPI (blue). Bar, 5 μm. (**B**) CaLS310 (*MCM2*/*MCM2*::*FRT*), where one copy of *MCM2* has been deleted, and two independent transformants, CaLS311 and CaLS312 (*MCM2*::*FRT*/*MET3prMCM2*) where the remaining wild-type copy was placed under the control of the *MET3* promoter of *C. albicans* were streaked on plates containing inducible (CM-met-cys) and repressible (CM+5 mM cys + 5 mM met) media and photographed after 48 h of incubation at 30°C. (**C**) Western blot of the whole cell lysate of CaLS306 (*MET3prMCM2/MCM2::FRT CSE4/CSE4-TAP*) using anti-Protein A antibodies shows time-dependant decrease in CENPA levels upon depletion of Mcm2 for 3, 6, 9 h. Normalization was performed using PSTAIRE. (**D**) CENPA cluster delocalized upon a prolonged depletion of Mcm2 in CaLS311 (*MET3prMCM2/MCM2*::*FRT CSE4-GFP-CSE4/CSE4*). Scale bar, 5 μm. (**E**) CENPA ChIP-qPCR using anti-protein A antibodies revealed significant reduction at *CEN7* in CaLS306 when grown either in CM-met-cys or CM+5mMmet+5mM cys for 6 h. (**F**) ChIP-qPCR in CaLS311 reveals no significant enrichment of Orc4 as compared to the permissive condition. Two-way ANOVA was used to determine statistical significance (***p<0.001, ns p>0.05).

In order to determine the essentiality of the *MCM2* gene in *C. albicans*, we constructed a conditional mutant of *mcm2* by deleting one allele and replacing the endogenous promoter of the remaining *MCM2* allele with the *MET3* promoter (Care, Trevethick et al. 1999). Mcm2 was found to be essential for viability in *C. albicans* (Figure 5B). Western blot analysis in the Mcm2 expressed versus depleted conditions confirmed the depletion of the protein levels of Mcm2 by 3 h (Figure S5E). We observed a drastic decline in the viability (Figure S5F) and an increased rate of mis-segregation of chromosomes in the *mcm2* mutant post 6 h of depletion (Figure S5G). We wanted to probe the cause of kinetochore mis-segregation by examining the effect of Mcm2 depletion on CENPA. Depletion of Mcm2 led to a concomitant reduction in CENPA protein levels (Figure 5C), similar to the previous observation on Orc4 depletion, shedding light on a previously unknown candidate to preserve kinetochore stability. We also observed declustering of the kinetochore architecture in the *mcm2* mutant (Figure 5D). ChIP-qPCR analysis revealed >50% reduction in the chromatin-bound CENPA following Mcm2 depletion for 6 h (Figure 5E). Strikingly, the centromeric occupancy of Orc4 was unaltered upon Mcm2 depletion for 6 h (Figure 5F). Hence, Mcm2 is required for CENPA stability but is dispensable for the centromeric binding of Orc4.

### The CENPA chaperone Scm3 loads CENPA during anaphase in *C. albicans*

Having identified previously unknown factors regulating CENPA stability, we wanted to examine the *de novo* loading of CENPA at the kinetochore in *C. albicans*. To address this, we sought to characterize the homolog of the chaperone Scm3/HJURP in *C. albicans*. BLAST analysis using *S. cerevisiae* Scm3 as the query sequence revealed that orf19.668 translates to a protein of ~72 kDa containing the CENPA-interacting Scm3 domain (Sanchez-Pulido, Pidoux et al. 2009) that was found to be conserved in *C. albicans* (Figure S6A). Additionally, there were three separate C2H2 zinc finger clusters present towards the C-terminus of Scm3 in *C. albicans* that was absent in *S. cerevisiae* (Aravind, Iyer et al. 2007) (Figure S6B). We generated a strain CaNV51 in which one of the endogenous copies of *SCM3* was tagged at the C-terminus with 2xGFP and a kinetochore protein Ndc80 was tagged with RFP. Microscopic examination revealed Scm3 signals as a distinct punctum in the nucleus colocalizing with Ndc80 at the anaphase stage of the *C. albicans* cell cycle (Figure 6A). Scm3 localization at other stages of the *C. albicans* cell cycle could not be detected. Also, we were unable to detect any signal in nocodazole treated cells of CaNV51 at metaphase (Figure 6A). This localization pattern of Scm3 coincides with the anaphase loading of CENPA in *C. albicans* (Shivaraju, Unruh et al. 2012) and hence, suggests that Scm3 indeed serves as the CENPA chaperone in *C. albicans*.

**Figure 6.**
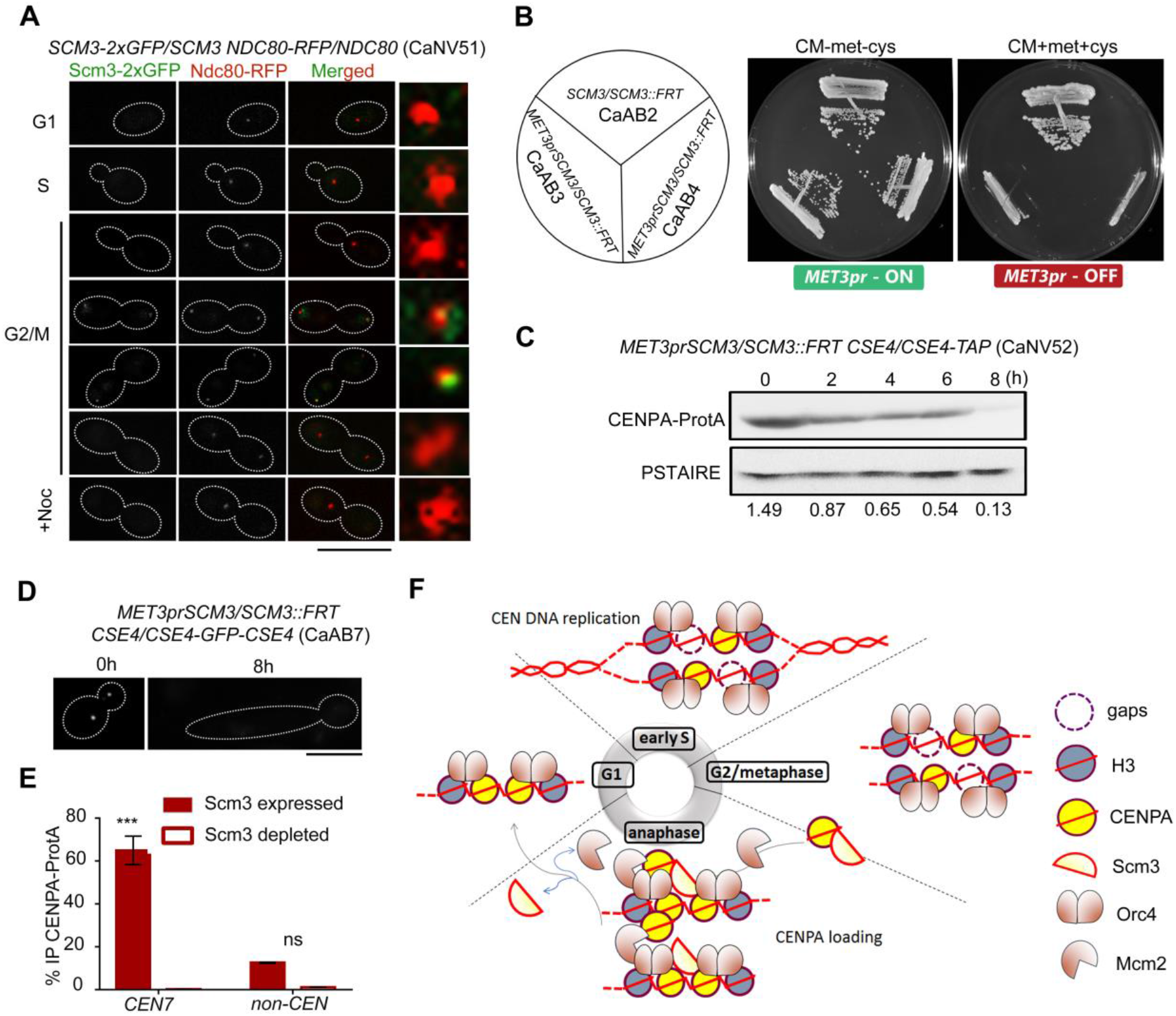
Scm3 is essential for the anaphase loading of CENPA in *C. albicans*. (**A**) Localization of Scm3 in CaNV51 (*SCM3-2xGFP/SCM3 NDC80-RFP/NDC80*) cells co-expressing Scm3-2xGFP and a kinetochore marker, Ndc80-RFP at various stages of the cell cycle. During anaphase, Scm3 colocalizes with the kinetochore cluster. Absence of Scm3 at the unsegregated kinetochore cluster in the budded CaNV51 cells upon treatment with 20 μg/ml nocodazole (NOC), last panel. The last panel displays zoomed-in view of the puncta. Bar, 10μm. (**B**) CaAB2 (*SCM3/SCM3*::*FRT*), where one copy of *SCM3* has been deleted, and two independent transformants CaAB3 and CaAB4 (*MET3prSCM3 /SCM3*::*FRT*), where the remaining allele was placed under the control of the *MET3* promoter were streaked on plates containing inducible (CM-met-cys) and repressible (CM+1 mM cys + 1 mM met) media and photographed after 48 h of incubation at 30°C. (**C**) Western blot showing protein levels of CENPA upon depletion of Scm3 for the indicated time points normalized to PSTAIRE. **(D)** Dissociation of the CENPA-GFP cluster in CaAB7 (*MET3prSCM3/SCM3::FRT CSE4/CSE4-GFP-CSE4*) cells grown in non-permissive conditions for the indicated time points. **(E)** CENPA ChIP-qPCR revealed significant reduction in the CENPA level at *CEN7* in CaNV52 (*MET3prSCM3/ SCM3::FRT CSE4/CSE4-TAP)* when grown in inducible (CM-met-cys) or repressible (CM+1mMmet+1mM cys) media for 8 h. (**F**) A model to explain CENPA loading at anaphase and CEN stabilization by the constitutive localization of Orc4 in *C. albicans*.

To determine the essentiality of Scm3, we constructed a depletion mutant of *scm3* by replacing the endogenous promoter of the intact copy with the *MET3* promoter in a heterozygous null strain (as described in previous sections) and found Scm3 to be essential for viability in *C. albicans* (Figure 6B). We observed the gradual degradation of CENPA protein levels by western blot following the gradual depletion of Scm3 (Figure 6C). Microscopic examination of CENPA revealed declustering of kinetochore post 8 h of Scm3 depletion, a phenotype observed upon depletion of any of the essential kinetochore proteins in *C. albicans* (Thakur and Sanyal 2012) (Figure 6D). Additionally, ChIP-qPCR analysis revealed a drastic reduction in CENPA from CENs upon Scm3 depletion (Figure 6E). Taken together, these results corroborate to the fact that Scm3 is localized to the kinetochore during anaphase and presumably loads and stabilizes CENPA as a CENPA-specific chaperone in *C. albicans*.

## Discussion

The organization of the genome into functional territories ensures complete and error-free DNA replication and faithful segregation of duplicated chromosomes. In the present study, we examine the genome-wide binding of Orc4 in *C. albicans*. Being strongly enriched at CENs, apart from other discrete genomic loci, Orc4 occupancy at CENs was strikingly similar to that of CENPA. Along the chromosomes, Orc4 was found to be associated with four distinct DNA binding modes, which are spatiotemporally positioned across the genome. The 3D genome model constructed in this study, which is the first to be performed in *C. albicans*, revealed that the early replicating highly enriched Orc4-bound regions are located more towards the clustered CENs and the mid- and late replicating regions are positioned towards the TELs. The centromeric localization of Orc4 was found to be constitutive and essential for CENPA stability, similar to Mcm2. We could also demonstrate the localization of Scm3 during anaphase suggesting that it is the CENPA-specific chaperone in *C. albicans*. Overall, we attempt to explain the mechanisms underlying CENPA establishment and stabilization in the presence of a constitutively bound protein, Orc4 at the centromeric chromatin.

Of the 414 genomic Orc4-bound regions identified in our study, 50 of them were located within the tRNA genes (tDNA). tDNAs contain binding sites for TFIIIC, TFIIIB, RNA polymerase III and SMC subunits (Glynn, Megee et al. 2004, Kogut, Wang et al. 2009) and exhibit a conserved replication timing (Muller and Nieduszynski 2017). tDNAs cluster near CENs and recover stalled forks (Thompson, Haeusler et al. 2003). The various Orc4 binding DNA motifs identified in our study hint towards differential usage and specification of replication origins facilitated by multiple modes of ORC binding in *C. albicans.* Such a differential mode of origin recognition has previously been demonstrated in *Pichia pastoris* which utilizes a combination of AT-rich and GC-rich origins to fulfil its genome duplication (Liachko, Youngblood et al. 2014). A previous genome-wide study on identification of ORC binding regions in *C. albicans* (Tsai, Baller et al. 2014) utilized antibodies against the *S. cerevisiae* ORC complex to report ~390 ORC binding sites, 25% (106/414) of which overlapped with our study. Since we used antibodies against an endogenous protein (CaOrc4) to map its binding sites in *C. albicans*, we possess a more authentic depiction of Orc4 occupancy in the genome.

In eukaryotes, early firing origins are more efficient and are organized into initiation zones (Mesner, Valsakumar et al. 2013). Our study shows this conserved feature in *C. albicans* as well, wherein orc^E^-orc^E^ and orc^L^-orc^L^ regions cluster significantly more than orc^E^-orc^L^ regions. The orc^E^ regions were found to be enriched with a higher ORC concentration than orc^L^ regions. One fact that could limit the resolution of our analysis is that the anti-Orc4 antibodies might primarily detect the early regions due to a higher enrichment of Orc4 at these sites, hence over-representing the “early” dataset. Similar to previous studies (Mesner, Valsakumar et al. 2011, Mesner, Valsakumar et al. 2013) early firing but not late firing origins appear to have been sequenced to saturation. In our analysis, we also find a higher contact probability between orc^L^-orc^L^ regions than orc^E^-orc^E^ regions. One explanation could be the fewer number of orc^L^ regions that we obtained from the tertile distribution. The orc^E^ regions form closely associated units and interact sparsely with orc^L^, reminiscent of the genome-wide replication landscape of a related species *Candida glabrata* (Descorps-Declere, Saguez et al. 2015). Hence, one can speculate the existence of topologically distinct domains that are separated in location and time as S phase progresses.

Over the years, there has been a more holistic understanding of ORC-origin recognition from DNA-based to chromatin-based to conformation based to the recently explored interactions mediated by multiprotein complexes that phase separate in solution. A recent evidence shows that ORCs are known to phase separate, which explains their strikingly non-uniform localization in the fly cell nucleus (Parker, Bell et al. 2019). ORCs in flies contain an intrinsically disordered region (IDR) that drives phase separation (Parker, Bell et al. 2019). Even though IDRs are absent in ORC subunits of yeast, the replication timing driven distribution of Orc4 identified in our study can help to ascertain conserved processes like replication origin communication, coordination of origin firing time and establishment and maintenance of heterochromatin. In *S. cerevisiae*, a ‘replication wave’ propagating from the centromeric regions enriched with early origins, through chromosomal arms and towards the late replicating sub-telomeric regions has been suggested (Lazar-Stefanita, Scolari et al. 2017). Polymer models for chromosomes previously generated using Hi-C contact maps assume an inverse relationship between contact probability and the average distance between the bead pairs/two loci. However, here we present a simulation method where the contact map is taken as input and the corresponding average 3D distance between any two regions in the genome is predicted. Our simulations produce an ensemble of steady-state genome configurations (corresponding to a population of cells) using which the 3D organization and other statistical properties of the genome can be computed. It would be of further interest to examine the 3D structure of the *C. albicans* genome in an *orc4* mutant.

The strong centromeric enrichment of Orc4 and its overlapping binding pattern with CENPA was particularly striking in our study because of the lack of a consensus DNA sequence at CEN in *C. albicans* (Sanyal, Baum et al. 2004). The high abundance of Orc4 at both the native CENs and a neocentromere strongly suggests its role in CEN establishment. The mode of Orc4 recognition at CENs is not DNA sequence-mediated as we could not detect any consensus among the four modes (A, B, C, D) across all eight CENs in *C. albicans*. However, we can speculate that the absence of H3 nucleosomes at CEN is a molecular cue for Orc4 binding and its subsequent stabilization. In humans, Orc2 has been shown to localize to CEN through its interaction with HP1 and is required for chromosome condensation and heterochromatin organization (Prasanth, Prasanth et al. 2004, Prasanth, Shen et al. 2010). *C. albicans* does not have conventional heterochromatin machinery (Freire-Beneitez, Price et al. 2016) nor do its CENs contain an active replication origin (Mitra, Gomez-Raja et al. 2014). In the context of the results obtained, the non-reciprocal relationship of Orc4 and CENPA proves that Orc4 is constitutively associated with centromeric chromatin and regulates CENPA stability. This is further supported upon Mcm2 depletion, which does not dislodge Orc4 from CEN, making Orc4 a component associated with kinetochore assembly. Whether Orc4 is directly bound to DNA or is recruited by a chromatin-mediated indirect interaction with the kinetochore needs to be tested in future. In humans, the S phase retention of CENPA is mitigated by its simultaneous interaction with HJURP and Mcm2 (Zasadzinska, Huang et al. 2018), which together transmit CENPA nucleosomes upon its disassembly ahead of the replication fork. In the light of the existing evidence in metazoan systems and the results obtained in our study, Mcm2 emerges as an evolutionarily conserved factor required for stabilizing CEN chromatin, perhaps through its transient interaction with Scm3. The anaphase specific loading of CENPA in *C. albicans* (Shivaraju, Unruh et al. 2012) which occurs much later than the S phase replenishment of ORCs, would also explain the non-reciprocal interdependency of Orc4 and CENPA, making Orc4 not only vital for CEN establishment but also for CEN maintenance in this organism.

Finally, we propose a model of CEN establishment and propagation in *C. albicans* (Figure 6F). Upon CEN replication during early S phase, CENPA is distributed equally into the daughter DNA molecules presumably leaving nucleosome-free regions. However, till the anaphase loading of CENPA by Scm3, these gaps have to be protected. In humans and flies, this process is mediated with the help of placeholder molecules like H3.3 (Dunleavy, Almouzni et al. 2011, Ray-Gallet, Woolfe et al. 2011). Since Orc4 is a constitutive component at the kinetochore, we propose it to be an effective stabilizer of centromeric chromatin till new CENPA is loaded during anaphase. This can be supported by the fact that ORCs have a propensity to bind to nucleosome depleted regions (Lipford and Bell 2001) and CENs are depleted of H3 nucleosomes. During anaphase, new CENPA is loaded by Scm3 and presumably through its transient interaction with Orc4 and Mcm2, the CENPA chaperone protects the CENPA nucleosomes. The acquisition of the novel module of three C2H2 domains in Scm3 of *C. albicans* which is absent in its *S*. *cerevisiae* counterpart might suggest a species-specific pathway for CENPA loading. It is to be noted that we could not detect the presence of Mcm2 at CENs when ChIP-sequencing was performed in asynchronous cells (data not shown). This could suggest a transient association of Mcm2 with CENs. Hence, we propose that Orc4 bookmarks CENs and in collaboration with Mcm2 and Scm3, in a manner analogous to human cells, directs CENPA to CEN DNA during anaphase in *C. albicans.*

## Materials and methods

### Construction of a conditional *orc4* mutant

In order to create a conditional null mutant of *orc4* in *C. albicans*, a deletion cassette was constructed as follows: a 368 bp fragment (Ca21Chr5 480170-479721) upstream of orf19.4221 was amplified using the primers ORC4_13/ORC4_14 from the genomic DNA of SC5314 and cloned as a KpnI/XhoI fragment into pSFS2a (Reuss, Vik et al. 2004) to create pLSK1. A 490 bp fragment (Ca21Chr5 478025-477535) downstream to orf19.4221was amplified using ORC4_15/ORC4_16 and cloned as a SacII/SacI fragment into pLSK1 to generate pLSK2. pLSK2 was linearized using KpnI and SacI, and used to transform YJB8675 (Joglekar, Bouck et al. 2008) and selected for nourseothricin resistance to obtain the strain CaLS328. The marker was recycled to obtain the nourseothricin sensitive strain CaLS329. To inactivate the remaining allele, a conditional mutant was constructed by cloning the N-terminus of orf19.4221 (Ca21Chr5 479720-479221) as a BamHI/PstI fragment in pCaDIS (Care, Trevethick et al. 1999). The resulting plasmid pLSK3 was linearized using BglII and transformed in CaLS329 to obtain independent transformants of the conditional mutant CaLS330, CaLS331. Similar deletions were performed in SN148 background to obtain the *orc4* conditional mutants CaLS322, CaLS323 and CaLS324. In each of these strains, a *CSE4-TAP-HIS* cassette (Mitra, Gomez-Raja et al. 2014) was transformed to obtain CaLS325, CaLS326 and CaL327, respectively. Transformants were confirmed by PCR and western blot analysis.

### Construction of a conditional *mcm2* mutant

In order to create a conditional null mutant of *mcm2* in *C. albicans*, a deletion cassette was constructed as follows: a 474 bp fragment (Ca21ChrR 857151-856675) upstream of orf19.4354 was amplified using the primers MCM2_13/MCM2_14 from the genomic DNA of SC5314 and cloned as a KpnI/XhoI fragment in pSFS2a to generate pLSK4. A 468 bp fragment (Ca21ChrR 853962-853494) downstream to orf19.4354 was amplified using MCM2_15/MCM2_16 and cloned as SacII/SacI fragment in pLSK4 to generate pLSK5. The plasmid was digested using KpnI and SacI, used to transform YJB8675 and selected for nourseothricin resistance to obtain the strain CaLS309. The marker was recycled to obtain the nourseothricin sensitive strain CaLS310. To inactivate the remaining allele, a conditional mutant was constructed by cloning the N-terminus of orf19.4354 (Ca21ChrR 856674-856164) as a BamHI/PstI fragment in pCaDIS (47). The resulting plasmid pLSK7 was linearized using BglII and used to transform CaLS310 to obtain independent transformants of the conditional mutant CaLS311, CaLS312 and CaLS313. Similar deletions were performed in SN148 background to obtain the *mcm2* conditional mutants CaLS303, CaLS304 and CaLS305. In each of these strains, a *CSE4-TAP-HIS* cassette (Mitra, Gomez-Raja et al. 2014) was transformed to obtain CaLS306, CaLS307 and CaLS308. Transformants were confirmed by PCR.

### Construction of a conditional *scm3* mutant

In order to create a conditional null mutant of *scm3* in *C. albicans*, a deletion cassette was constructed as follows: a 598 bp fragment (Ca21Chr3 390264-390708) upstream of orf19.1668 was amplified using the primers ASB25/ASB26 from the genomic DNA of SC5314 and cloned as a KpnI/XhoI fragment in pSFS2a to create pASB1. A 305 bp fragment (Ca21Chr3 387626-388030) downstream to orf19.1668 was amplified using ASB27/ASB28 and cloned as SacII/SacI fragment in pASB1 to generate pASB2. The plasmid was digested using KpnI and SacI, used to transform *C. albicans* SN148 and selected for nourseothricin resistance to obtain the strain CaASB1. The marker was recycled to obtain the nourseothricin sensitive strain CaAB2. To inactivate the remaining allele, a conditional mutant was constructed by cloning the N-terminus of orf19.1668 (Ca21Chr3 389106-390089) as a BamHI/PstI fragment in pCaDIS (47). The resulting plasmid pAB3 was linearized using KpnI and transformed in CaAB2 to obtain independent transformants of the conditional mutant CaAB3. The conditional mutants were further confirmed by PCR. Similar deletions were performed in YJB8675 and CaKS102 backgrounds. To construct the *MET3*pr*-SCM3* cassette with a *HIS1* marker, a *HIS1* fragment was cloned into the EcoRI site of the plasmid pASB3 to obtain pASB4. The plasmid pAB4 was linearized with KpnI and was used to transform CaAB9 to obtain CaNV52. The conditional mutants were confirmed by their inability to grow in non-permissive media.

### Construction of a strain expressing a Protein A tag at the C-terminus of Mcm2

The strain CaKS107 (*MCM2/MCM2-TAP*) was constructed by integrating a C-terminal Prot A tagging cassette with NAT marker created by overlap extension PCR using the primers M1 to M6 in BWP17. To delete the remaining allele of *MCM2*, the deletion cassette pLSK5 was modified as follows: The *NAT* marker from pLSK5 was released using BamHI/PstI and a *URA3* fragment digested with the same enzymes was cloned into this backbone to generate pLSK6. This plasmid was digested using KpnI/SacI and transformed into CaKS107 to generate CaLS334 (*MCM2-TAP(NAT))/URA3*). *URA3* was recycled from this strain to obtain CaLS335, CaLS336 and CaLS337. The deletion was confirmed by PCR with specific primers and expression of the tagged protein was confirmed with western blot using anti-Protein A antibodies. To replace the endogenous promoter of *MCM2* with the *MET3* promoter, pLSK7 was transformed into CaLS335, CaLS336 and CaLS337 to generate CaLS338, CaLS339 and CaLS340.

### Construction of a strain expressing 2xGFP epitope-tagged Scm3 under the native promoter and Ndc80 tagged with RFP

To study the subcellular localization of Scm3 under the native promoter, we constructed the strain expressing Scm3-tagged with a double GFP at its C-terminus under the native promoter in the strain SN148. The 3’coding sequence of *SCM3* excluding the stop codon was amplified with the oligos NV158 and NV159 and cloned into the SacII and SpeI sites of pBSGFP-URA3, to obtain pNV31. Another fragment of the *GFP* ORF was amplified using oligos NV250 and SR67 and inserted into the SpeI site of pNV31 to obtain pNV32. After confirming the orientation of GFP by HpaI, the plasmid was linearized with SwaI and was used to transform to obtain CaNV50. Further, to simultaneously localize Scm3 and Ndc80, we transformed CaNV50 with pNdc80-RFP-ARG4 (Varshney and Sanyal 2019) after linearizing with XhoI to obtain CaNV51. The transformants were screened by microscopy.

### Generation of anti-Orc4 antibodies

The peptide sequence from *C. albicans* Orc4 (YLPKRKIDKEESSI) was chemically synthesized and conjugated with Keyhole Limpet Hemocyanin. The conjugated peptide (1 mg/ml) was mixed with equal volumes of Freund’s complete adjuvant and used as an antigen to inject non-immunized rabbits as the priming dose. Three subsequent booster doses at an interval of two weeks (per immunization) were given using Freund’s incomplete adjuvant. Following antibody detection using ELISA, major bleed was performed. The anti-serum was collected, IgG fractionated and affinity purified against the free peptide (AbGenex, India). The specificity of the purified antibody preparation was confirmed by western blot and immunolocalization experiments.

## Method details

### Media and growth conditions

*ORC4* and *MCM2* mutants were grown either in CM-methionine-cysteine or in CM + 5mM methionine +5mM cysteine for the indicated number of hours. *SCM3* mutants were grown in presence of 1 mM methionine+ 1mM cysteine for repression. CAKS3b (Sanyal and Carbon 2002) was grown in YP with succinate (2%) for expressing CENPA and YP with dextrose (2%) for depleting CENPA for 8 h for the ChIP experiments. SC5314 was grown in YPDU. The *cdc15* mutant SBC189 (Bates 2018) was grown in CM and repressed in presence of 20 μg/ ml doxycycline for 16 h. To arrest cells in the S phase, YJB8675 (Joglekar, Bouck et al. 2008) were grown in presence of 200 mM hydroxyurea for 2h. To arrest cells in metaphase, YJB8675 and CaNV51 cells were grown in presence of 20 μg/ ml nocodazole for 4 h.

### Western blotting

Approximately 3 O.D. equivalent cells were harvested and precipitated by 12.5% TCA overnight at −20°C. The pellet was spun down at 13000 rpm and washed with 80% acetone. The pellet obtained was then dried and resuspended in lysis buffer (1% SDS, 1N NaOH) and SDS loading dye. Samples were boiled for 5 min and electrophoresed on a 10% polyacrylamide gel. Protein transfer was performed by semi-dry method for 30 min at 25V. Following protein transfer, the blot was blocked with 5% skimmed milk for an hour. The blot was incubated with primary antibodies in the following dilutions: rabbit anti-Protein A (1:5000)/ rabbit anti-Orc4 (1:1,000)/ mouse anti-PSTAIRE (1: 5000). The blot was washed thrice in PBST (1X PBS + 0.05% Tween) and incubated with goat anti-rabbit IgG-HRP (1:10,000) or goat anti-mouse IgG-HRP (1:10,000). Following three PBST washes, the blot was developed using chemi-luminescence method. For quantifying protein level with respect to PSTAIRE, band intensity of the desired protein was divided with that of PSTAIR in the corresponding lane and ratio was calculated using densitometric analysis.

### Indirect immuno-fluorescence

Exponentially grown cultures of SC5314 and CaLS335 were fixed with 37% formaldehyde. Spheroplasts were prepared using lysing enzyme and cells were fixed on poly-lysine coated slides using methanol and acetone and then incubated with 2% skimmed milk to block non-specific binding. Following ten PBS washes, cells were incubated with anti-Orc4 antibodies (1:100) or anti-Protein A antibodies (1:1,000) for 1 h in a humid chamber. Post PBS washing, cells were incubated with the Alexa Fluor goat anti-rabbit IgG 568 (1:500) or Alexa Fluor goat anti-rabbit IgG 488 (1:500) for 1 h. The slide was mounted on a coverslip using DAPI (10 ng/ul). Microscopic images were captured by a laser confocal microscope (Carl Zeiss, Germany) using LSM 510 META software with He/Ne laser (bandpass 565-615 nm) for Alexafluor 568 and a 2-photon laser near IR (bandpass~780 nm) for DAPI. Z-stacks were collected at 0.4-0.5 μm intervals and stacked projection images were processed in ImageJ and Adobe Photoshop.

### Live cell microscopy

For conditional expression of genes under the *MET3* promoter, GFP-tagged strains were grown in permissive media (CM -met-cys) overnight and repressed in presence of CM + 5 mM met+5mM cys or CM + 1 mM met+1 mM cys for the indicated time. In each case, the cells were washed twice with water and resuspended in distilled water which was placed on a 2% agarose bed on a glass slide.

DeltaVision System (Applied Precision) was used for CENPA localisation upon Scm3 depletion and Axio Observer Calibri (ZEISS) was used for localization of Scm3 and CENPA upon Mcm2 and Orc4 depletion. Images were processed using ImageJ and Adobe Photoshop. For quantification of relative GFP intensity, pixel values (arbitrary units) from the background fluorescence was subtracted from the pixel values obtained from the CENPA cluster (GFP) from individual cells. This was performed for 20 different small-budded cells of the control population and 20 cells arrested at S phase upon HU treatment and plotted in a scatter plot with standard error of mean (SEM). Students’ unpaired t-test was used to determine statistical significance.

### Chromatin Immunoprecipitation (ChIP)

For the Orc4 ChIP-sequencing experiment, approximately 500 O.D. of asynchronously grown log phase culture of *C. albicans* SC5314 cells was crosslinked for 1 hr using formaldehyde at a final concentration of 1%. The quenched cells were incubated in a reducing environment in presence of 9.5 ml distilled water and 0.5 ml of beta mercapto-ethanol. The protocol for ChIP was followed as described previously (Yadav, Sun et al. 2018). Briefly, the sheared chromatin was split in two fractions, one of which was incubated with 5ug/ ml IP of purified anti-Orc4 antibodies. Following overnight incubation, the IP and mock (no antibody) fractions were further incubated with Protein A-Sepharose beads. The de-crosslinked chromatin was purified and ethanol precipitated. The DNA pellet was finally resuspended in 20 μl of MilliQ water. All three samples (I, +, −) were subjected to PCR reactions. For the CENPA ChIP, cells were crosslinked for 15 min with 1% formaldehyde and IP samples were incubated with 4 μg/ ml of anti-Protein A antibodies or 3 μg/ml of anti-GFP antibodies. Rest of the protocol was the same as described above.

### ChIP-sequencing analysis

Immunoprecipitated DNA and the corresponding DNA from whole cell extracts were quantified using Qubit before proceeding for library preparation. Around 5 ng ChIP and total DNA were used to prepare sequencing libraries using NEBNext Ultra DNA library preparation kit for Illumina (NEB, USA). The library quality and quantity were checked using Qubit HS DNA (Thermo Fisher Scientific, USA) and Bioanalyzer DNA high sensitivity kits (Agilent Technologies, USA) respectively. The QC passed libraries were sequenced on Illumina HiSeq 2500 (Illumina Inc., USA). HiSeq rapid cluster and SBS kits v2 were to generate 50 bp single end reads. The reads were aligned onto the *Candida albicans* SC5314 reference genome (v. 21) using bowtie2 (v. 2.3.2) aligner (Langmead, Trapnell et al. 2009). More than 95% of the reads mapped onto the reference genome (Control:97.74%; IP:96.13%). The alignment files (BAM) were processed to remove PCR duplicate reads using Mark Duplicates module of Picard tools. These processed BAM files were further taken for identification of peaks by MACS2 (Feng, Liu et al. 2012). These peaks were annotated with the *C. albicans* SC5314 reference annotation file. Visualisation of the aligned reads (BAM files) on the reference genome was performed using Integrative Genome Viewer (IGV) (https://software.broadinstitute.org/software/igv/) (Robinson, Thorvaldsdottir et al. 2011).

### ChIP-qPCR analysis

All ChIP-qPCR experiments were performed in three independent transformants with three technical replicates for each transformant. The input and IP DNA were diluted appropriately and qPCR reactions were set up using specific primers. ChIP-qPCR enrichment was determined by the percentage input method. In brief, the Ct values for input were corrected for the dilution factor (adjusted value= Ct value of input or IP minus log2 of dilution factor) and then the percent of the input chromatin immunoprecipitated by the antibody was calculated using the formula:100*2^ (adjusted Ct input-adjusted Ct of IP) (Mukhopadhyay, Deplancke et al. 2008). One-way ANOVA, two-way ANOVA and Bonferroni post-tests were performed to determine statistical significance. For all the Orc4 ChIPs, percent IP values at the CEN were either compared with the control region, *LEU2* or normalized with percentage IP values at a far-CEN Orc4 binding region.

### Hi-C analysis

Wild-type *C. albicans* Hi-C data was downloaded from PRJNA308106 (Burrack, Hutton et al. 2016). To examine interactions between the Orc4 binding regions, Hi-C interactions were analyzed according to the chromosome coordinates, different modes identified by DIVERSITY and also based on replication timing (orc^E^, orc^M^ and orc^L^). The heatmap for the full genome was plotted using log-scaled values with a pseudocount of 0.000001 (10^−6^). The heatmap for the “ORC-only” was plotted using values for the 2 kb windows overlapping with the midpoints of the Orc4 binding regions, using the same scaling and colour scale as the full-genome heatmap. The violin plots were calculated for 1,000 randomizations of each dataset, where for each randomization, the chromosomal distribution and lengths of the regions were preserved.

### Motif analysis by DIVERSITY

For motif analysis, the *de novo* motif discovery tool DIVERSITY (Mitra, Biswas et al. 2018) was used with default web-server options on the 417 Orc4 ChIP-seq peaks. DIVERSITY is specially developed for ChIP-seq experiments profiling proteins that may bind DNA in more than one way.

### Replication timing analysis

To analyze the replication timing of the ORC binding regions, fully processed timing data available in GSE17963_final_data.txt (Koren, Tsai et al. 2010) was used. A larger replication time value implies earlier replication. All the 414 genomic origins were aligned according to their timing scores, and categorized as 218 early, 127 mid and 69 late replicating regions based on the tertile distribution.

### Polymer modelling of chromosomes

The paired-end reads from the Hi-C data (Burrack, Hutton et al. 2016) were mapped onto the wildtype *C. albicans* genome assembly 21 following the Hi-CUP pipeline with default parameters (Wingett, Ewels et al. 2015). Next, the resulting bam file was analyzed using dryhic R package (Vidal, le Dily et al. 2018) and ICE normalization was applied. The contact matrix was converted to a data frame object and written to a file for subsequent analysis. To compute the 3D organization of the *C. albicans* genome, the chromatin was modelled as a polymer with N beads connected by (N – 1) harmonic springs. We coarse-grained the chromatin into equal-sized beads, each representing 10 kb of the genome and the connecting springs with a natural length l_0_ (Lieberman-Aiden, van Berkum et al. 2009). To model the haploid yeast genome of *C. albicans* comprising of 8 chromosomes of different lengths, we considered 8 polymer chains each consisting of 319, 224, 180, 161, 120, 104, 95 and 229 beads, respectively. The bead corresponding to the midpoint of CEN of each chromosome was assigned as the CEN bead. In each chain, to represent the connectivity, all neighbouring beads were connected linearly by a harmonic spring having energy (Ganai, Sengupta et al. 2014)

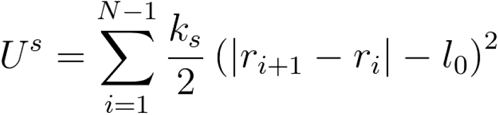

where *U*^s^ is the spring potential energy, k_s_ is the spring stiffness, r_i_ is the position vector of the i^th^ bead, and l_0_ is the natural length. The summation here is between nearest neighbours. To mimic the steric hindrance between any two parts of chromatin, the repulsive part of the Lennard-Jones (LJ) potential energy, given below, was used:

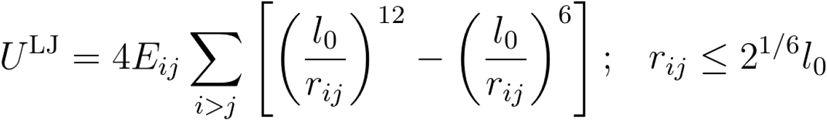

where r_ij_ represents the distance between bead i and bead j, *E*_ij_ represent the strength of attraction and the sum is over all possible bead-pairs. Hi-C data at a 10 kb resolution was considered as an input in the current model. We generated an initial configuration by connecting each pair of beads (i,j) with probability P_ij_ as per the Hi-C contact matrix. We chose a uniformly distributed random number r in the interval [0, 1] and the bond was introduced if P_ij_>r, for each pair of beads. This bond is also a harmonic spring with high stiffness kc and of natural length l0 having energy

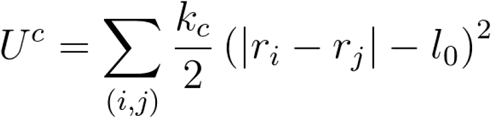

where (i,j) represent summation over the pairs< selected probabilistically as described above. All the chromosomes are confined into a sphere of radius R_s_ which represents the confinement arising due to the nucleus. One of the CENs (*CEN1*) was tethered to the nuclear periphery. The resulting polymer was equilibrated via Langevin simulation using LAMMPS (Plimpton 1995). The whole process described above was repeated for 1,000 realizations generating an ensemble of 1,000 configurations. Each of the configurations is equivalent of chromatin in a single cell.

## Accession number

ChIP-sequencing data used in the study have been submitted to NCBI under the BioProject accession number PRJNA477284.

## Acknowledgments

We thank Clevergene Biocorp for ChIP-seq experiments and analysis. We also thank Prakash for animal facility and B. Suma for confocal microscopy, JNCASR. We thank A. Koren and J. Berman for sharing the raw data of the replication timing experiment. We thank S. Bates, University of Exeter, UK for sharing the *cdc15* mutant strain and guiding us with the depletion protocol. We thank S. Mitra for constructing CaKS107 strain. We also thank R. Dighe, IISc, Bangalore for helping us in raising polyclonal antibodies. LS thanks support from Council of Scientific and Industrial Research (CSIR), Govt. of India grant number 09/733(0178)/2012-EMR-I and intramural support from Jawaharlal Nehru Centre for Advanced Scientific Research, Bangalore (JNCASR). KK and RP acknowledge support from SERB and DST India grant EMR/2016/005965. AB is supported by the grant BT/PR14840/BRB/10/880/2010. NV is supported by CSIR fellowships 09/733 (0253)/219-EMR-I and 9/733 (0161)/2011-EMR-I. BT thanks intramural support from JNCASR. KG acknowledges CSIR SPM fellowship (SPM-07/733(0181)/2013-EMR1) and financial support from JNCASR. LN is supported by DBT grant BT/ PR16240/BID/7/575/2016 and RS thanks the PRISM-II project at IMSc, funded by DAE. This project is also funded by Tata Innovation Fellowship, Dept. of Biotechnology, Govt of India to KS. KS also gratefully acknowledges the intramural funding from JNCASR.

## Author contributions

KS supervised the study. KS and LS conceived the idea and designed the experiments. LS constructed strains and reagents, performed and analyzed experiments pertaining to Orc4 and Mcm2. KK and RP simulated the chromosome model. Scm3 characterization and western blotting experiments were performed by AB; Scm3 localization and ChIP experiments were performed by NV. LN performed the motif analysis. RS and LN performed the Hi-C and replication timing analyses. BT performed the ChIP-sequencing analysis. KG analyzed the Hi-C data and derived the contact matrix. LS and KS wrote the manuscript with support from all of the authors. KS edited the manuscript and provided the funding.

