## Supplemental Information for "Orc4 spatiotemporally stabilizes centromeric chromatin"

### SUPPLEMENTAL FIGURES

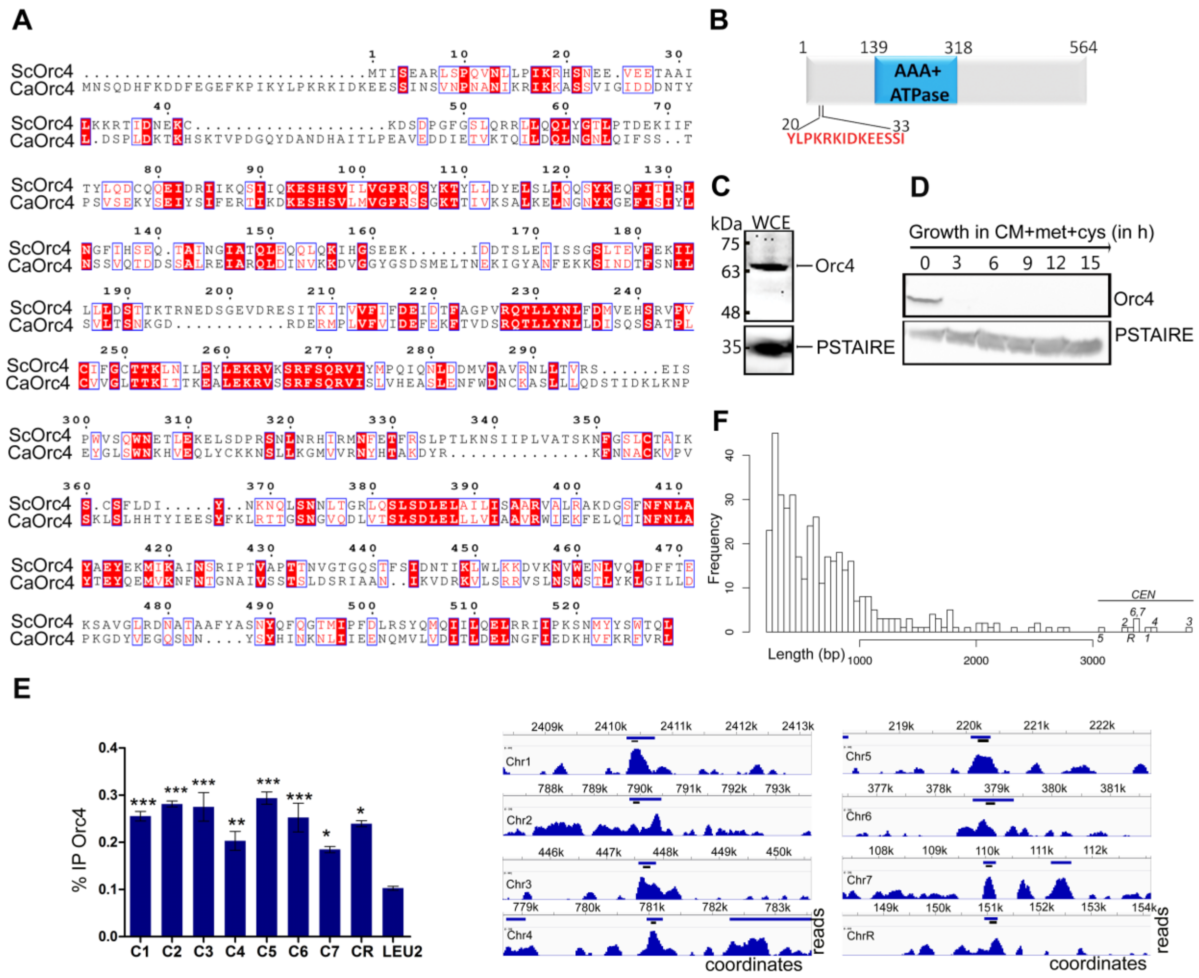

**Figure S1** (related to Figure 1). **Expression and *in vivo* localization of Orc4 in *C. albicans*.** (**A**) A comparison of the Orc4 proteins from *S. cerevisiae* and *C. albicans* reveals conserved stretches of amino acids constituting the AAA+ ATPase domain (aa 139-318). (**B**) Domain architecture of Orc4 reveals a 564 aa long polypeptide consisting of a central AAA+ ATPase domain in *C. albicans*. The peptide sequence chosen to raise the antibodies has been highlighted in red letters (aa 20-33). (**C**) Expression of Orc4 was verified by western blot with anti-Orc4 antibodies using whole cell extract (WCE) from *C. albicans* SC5314. Anti-Orc4 antibodies yielded a band at the expected molecular weight at ~64 kDa in a denaturing SDS PAGE. PSTAIRE was used as the loading control. (**D**) Western blot analysis using anti-Orc4 antibodies indicates time course depletion of Orc4 in the conditional mutant CaLS330 when the strain was grown for the indicated time (0, 3, 6, 9, 12, 15 h) in presence of 5 mM met + 5 mM cys. PSTAIRE was used as the loading control. (**E**) Orc4 ChIP followed by standard qPCR assays were used to validate the binding of Orc4 on one region in each of the eight *C. albicans* chromosomes (*left*). Primers (see Key Resource Table) corresponding to an Orc4-enriched region on each of chromosome 1 - 7 (C1- C7) and chromosome R (CR) were used to amplify the Orc4 ChIP DNA (*right*). Black lines indicate position of the amplicon for each of the C1-C7, CR loci. Each of the test regions were compared with *LEU2* to determine statistical significance using one-way ANOVA followed by Bonferroni post-tests (*** p<0.001, ** p<0.01, * p<0.05 ns: p>0.05). (**F**) Frequency distribution of the length of Orc4 binding sites in the *C. albicans* genome.

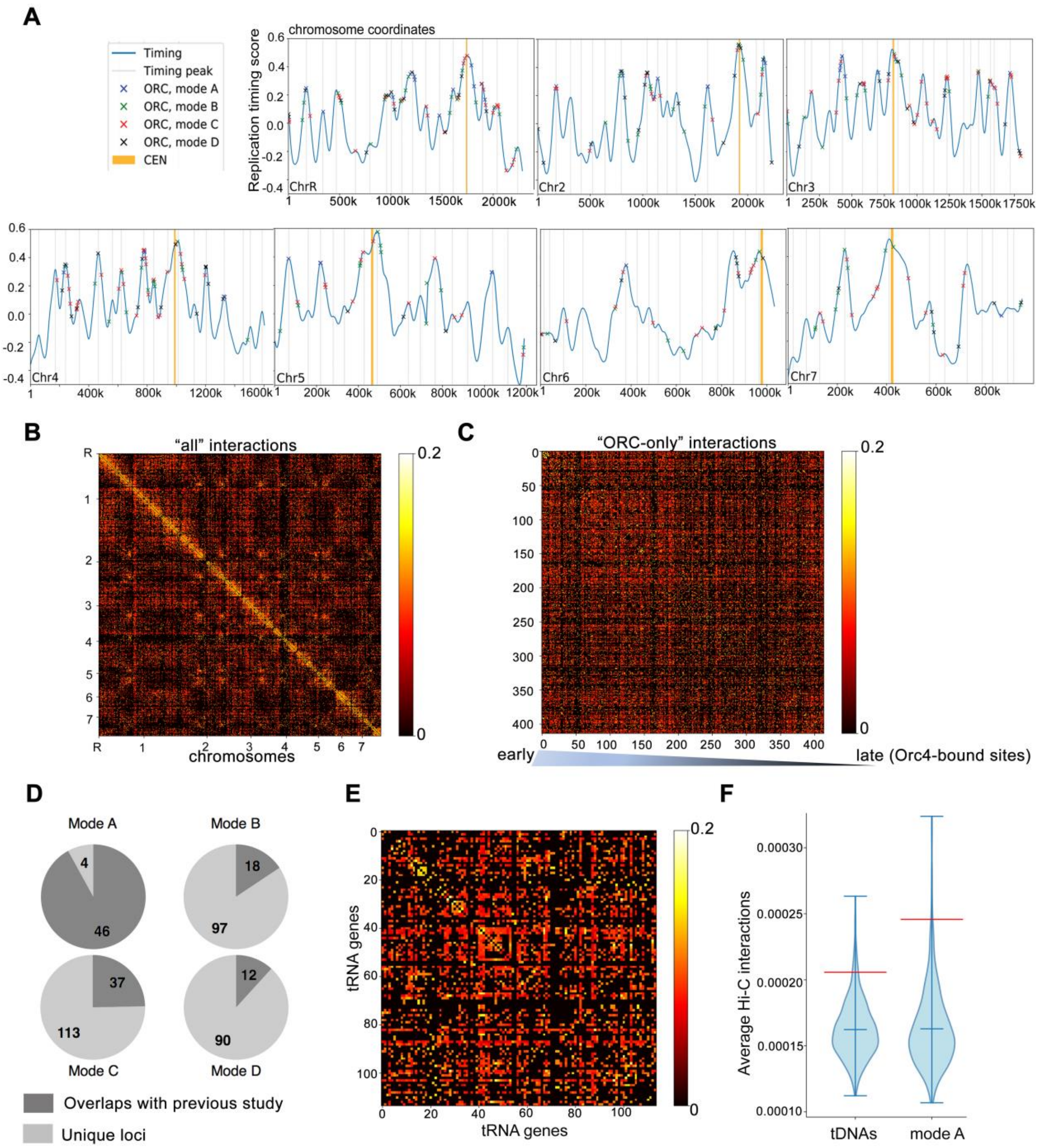

**Figure S2** (related to Figure 2). **Early replicating regions interact among themselves to form clusters/ replication domains.** (**A**) Orc4 ChIP-seq peaks aligned to the replication timing profile of *C. albicans* chromosomes from the previous study (Koren, Tsai et al. 2010). (**B**) The Hi-C heatmap shows a whole-genome "all" heatmap representation of the Hi-C data (Burrack, Hutton et al. 2016) as a 7145x7145 matrix with a 2 kb resolution. The maximum value in the data was 0.2015 and the minimum was zero. For plotting, the values were log-transformed with a pseudocount of 0.0001. (**C**) The Hi-C "ORC-only" heatmap shows interactions between the 414 chromosomal ORC binding regions, ordered by timing (early to late), to the same color scale as in (A). The analysis was performed at a resolution of 2 kb. It indicates marginally higher interactions within orc$^{E}$ (top-left quadrant) and within orc$^{L}$ (square on bottom right), in agreement with Figure 2C. (**D**) Pie charts depicting the number of Orc4 binding sites in the current study overlapping with binding sites reported earlier (Tsai, Baller et al. 2014). (**E**) Hi-C interactions for 50 tRNA-associated Orc4 peaks

(0-49) and 64 tRNA genes not overlapping with Orc4 peaks (50-113). (**F**) Average Hi-C interactions compared to 1,000 randomizations for the 64 non-Orc4 tRNA genes and the Orc4 peaks, suggesting tRNA genes in general interact more than average, but less than the tRNA-associated Orc4 peaks.

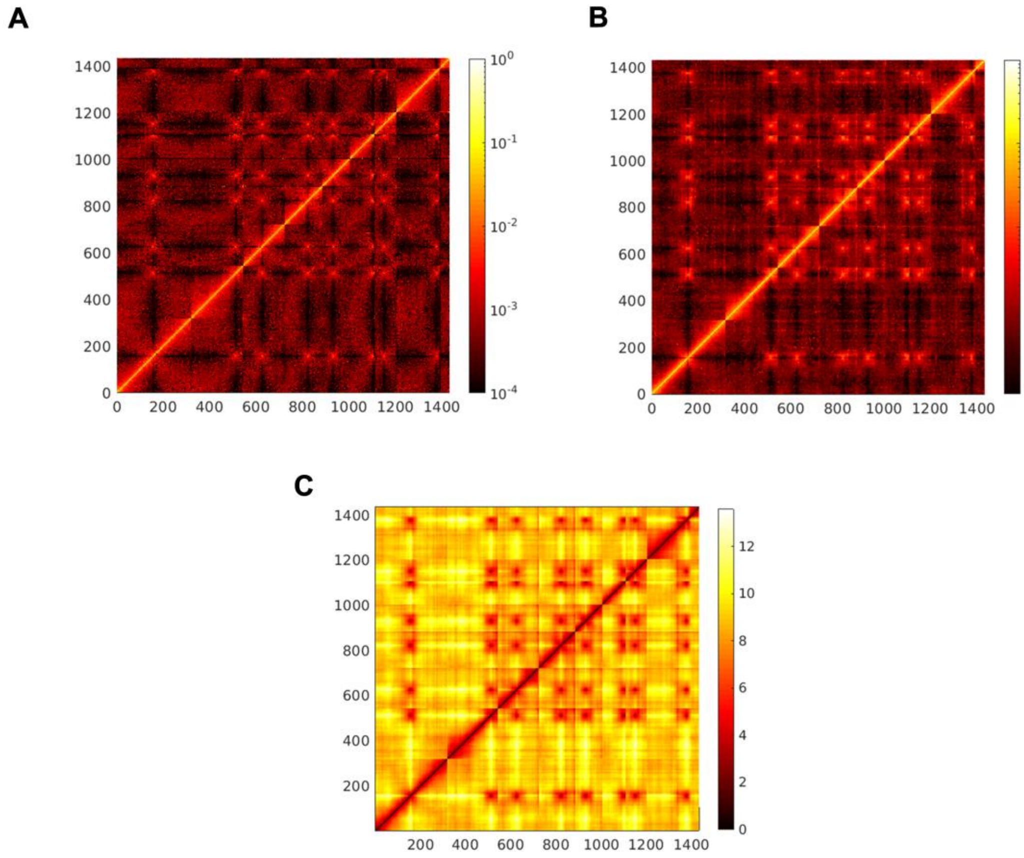

**Figure S3** (related to Figure 3). **Comparison between experimental and simulated Hi-C contact probabilities in *C. albicans***. (**A**) The Hi-C contact probability from the previous data (Burrack, Hutton et al. 2016) at a resolution of 10 kb and (**B**) contact probability calculated from the simulation over 1,000 realizations at a resolution of 10 kb. Here the abscissa and ordinate represent the bead number along the polymer chain, and gradient of the red color (bright to dark) indicate the contact probability between the bead pairs from high to low (see color bar). (**C**) Average spatial distances computed from the simulation based on the 1,000 independent configurations generated. Here the abscissa and ordinate represent the bead number along the polymer chain and gradient of color (yellow to red) indicate the spatial distance between the bead pairs from high to low.

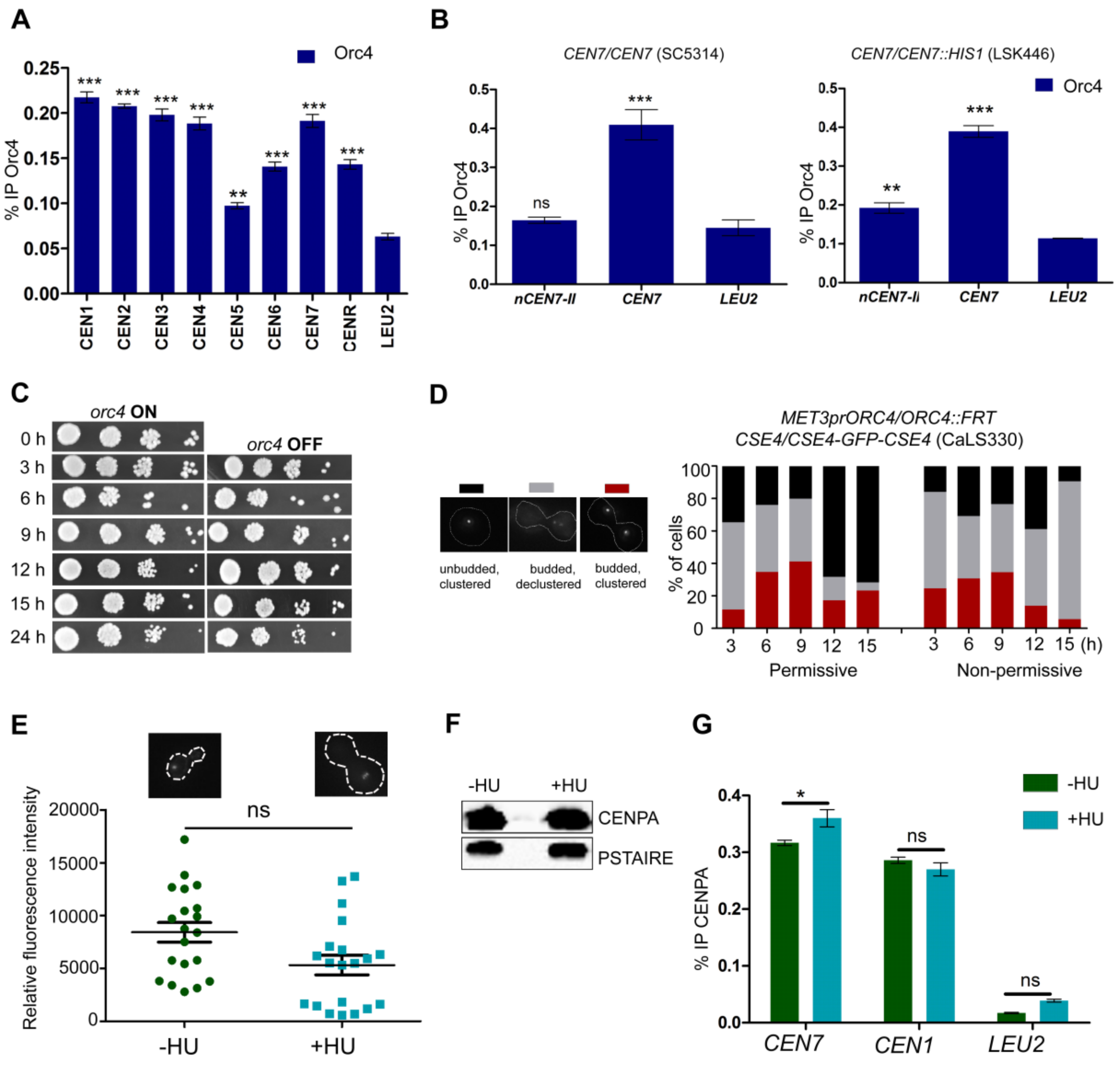

**Figure S4** (related to Figure 4). **Orc4 binds to active centromeres and facilitates chromosome segregation in *C. albicans*.** (**A**) Orc4 ChIP followed by standard qPCR assays in SC5314 was used to validate the enrichment of Orc4 at all *C. albicans* centromeres. *LEU2* was used as a control region. Statistical significance was determined by one-way ANOVA followed by Bonferroni post-tests (*** p<0.001, ** p<0.01, ns: p>0.05). (**B**) Orc4 ChIP qPCR in the wild type (*CEN7/CEN7*) (*left*) and *CEN7* deletion strain LSK446 (*CEN7/CEN7::HIS1)* (*right*) indicates significant enrichment of Orc4 at *nCEN7-II,* the neocentromere hotspot, over the control region (*LEU2*). Statistical significance was determined by one-way ANOVA followed by Bonferroni post-tests (*** p<0.001, ** p<0.01, ns: p>0.05). (**C**) Spot dilution assays to indicate viability in the *orc4* conditional mutant CaLS330 (*MET3prORC4/ORC4::FRT*) grown in permissive (CM-met-cys) or non-permissive media (CM+met +cys) for the indicated time and then spotted on CM-met-cys media. Plate photographs were captured after 48 h of incubation at 30ºC. (**D**) The segregation pattern of the clustered kinetochores (CENPA signals) was examined in CaLS330, an *orc4* conditional mutant when grown in permissive or non-permissive media for 3,6,9,12 and 15 h. The percentage of cells showing a specific segregation phenotype of clustered kinetochores in unbudded (black), unsegregated budded (grey) and segregated budded (red) was counted. At least 100 cells from three independent transformants of *orc4* mutant were analyzed for this assay. (**E**) Normalized fluorescence intensity of cells expressing CENPA-GFP when treated with or without 200 mM HU (n=20). Students unpaired t-test was used to determine statistical significance. (**F**) Western blot analysis using anti-Protein A antibodies in the strain J121 (*CSE4/CSE4-TAP(NAT)*) (Thakur and Sanyal 2013) reveals comparable levels of CENPA before (-HU) and after HU treatment (+HU). PSTAIRE was used as a loading control. (**G**) ChIP-qPCR analysis using anti-GFP antibodies reveals no depletion of CENPA at CENs upon HU treatment. Statistical significance was determined using two-way ANOVA (* p<0.05, ns: p>0.05).

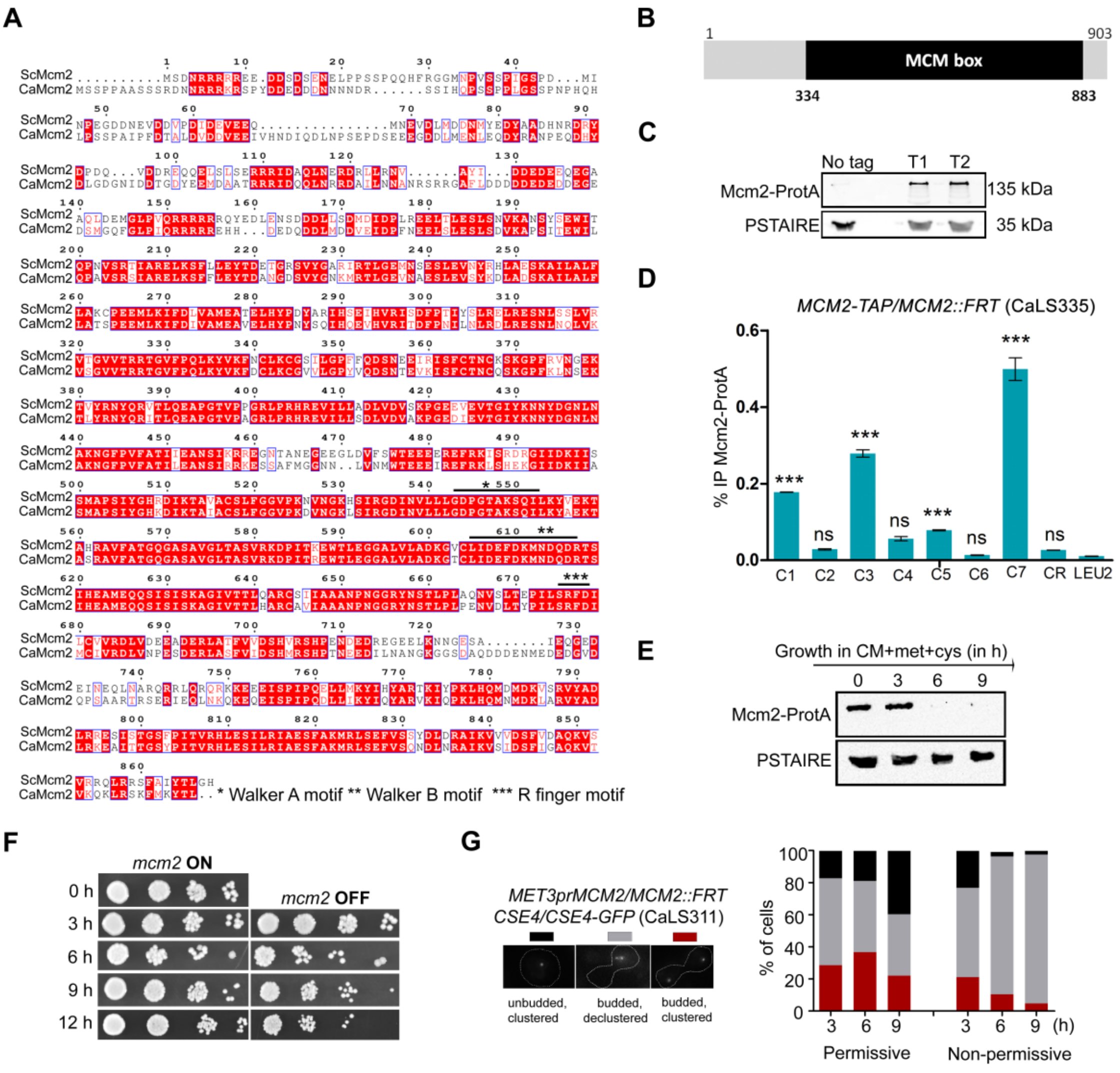

**Figure S5** (related to Figure 5). **Mcm2 is an evolutionarily conserved protein and is important for chromosome segregation in *C. albicans*** (**A**) A comparison of the Mcm2 proteins from *S. cerevisiae* and *C. albicans* reveals the three conserved motifs, Walker A, Walker B and R finger motif. (**B**) Domain architecture of Mcm2 reveals a 903 aa long polypeptide consisting of the catalytic helicase domain, MCM box (aa 334-883). (**C**) Western blot analysis of the whole cell extracts of CaLS335 *(MCM2-TAP(NAT)/MCM2::FRT)* using anti-Protein A antibodies displays a specific band of Mcm2-TAP corresponding to 135 kDa, which is undetected in the untagged strain BWP17. PSTAIRE was used as the loading control. (**D**) ChIP-qPCR analysis to examine the binding of Mcm2 to Orc4-bound regions on all eight chromosomes in *C. albicans* compared to a control region (*LEU2*). One-way ANOVA was used to determine statistical significance. (*** p<0.001, ns: p>0.05). (**E**) Western blot analysis using anti-ProtA antibodies indicates time course depletion of CENPA in CaLS306 (*MET3prMCM2-TAP(NAT)/MCM2::FRT*) when the strain was grown for the indicated time (0, 3, 6, 9h) in presence of 5 mM met +5 mM cys. PSTAIRE was used as the loading control. (**F**) Spot dilution assays to indicate viability in the *mcm2* conditional mutant CaLS311 (*MET3prMCM2/MCM2::FRT*) grown in permissive (CM-met-cys) or non-permissive media (CM+met +cys) for the indicated time and then spotted on CM-met-cys media. Plate photographs were captured after 48 h of incubation at 30ºC. (**G**) The segregation pattern of the clustered kinetochores (CENPA signals) was examined in CaLS311 when grown in permissive or non-permissive media for 3,6 and 9 h. The percentage of cells showing a specific segregation phenotype of clustered kinetochores in unbudded (black), unsegregated budded (grey) and segregated budded (red) was counted. At least 100 cells from three independent transformants of *mcm2* mutant were analyzed for this assay.

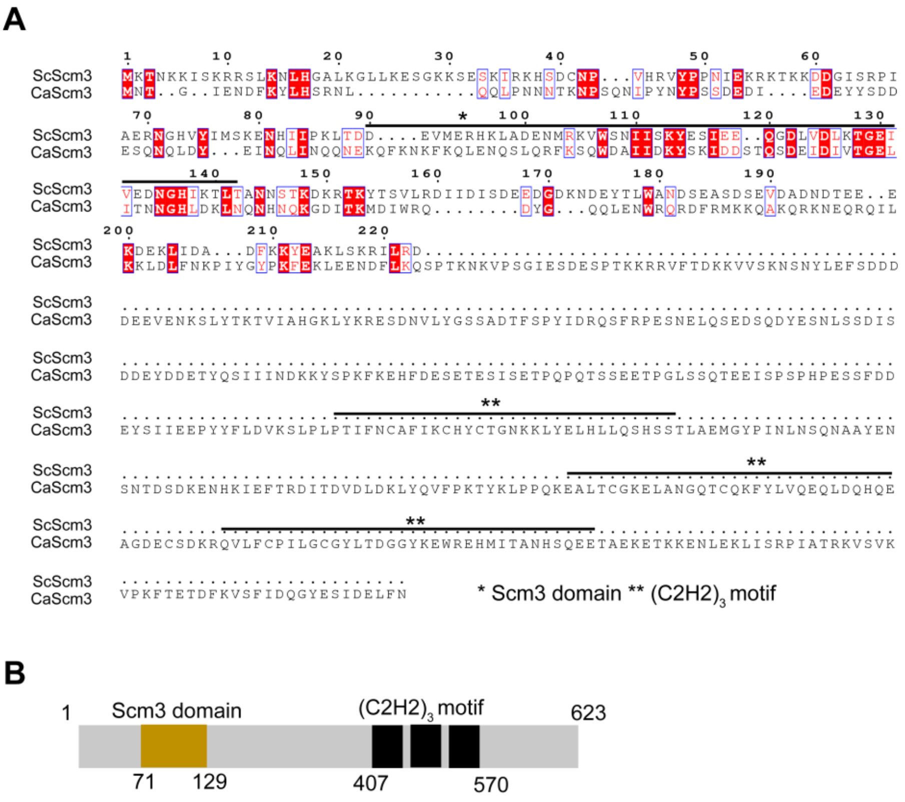

**Figure S6** (related to Figure 6). **Scm3 is a conserved protein in *C. albicans* (A)** A comparison of the Scm3 proteins from *S. cerevisiae* and *C. albicans* the conserved Scm3 domain (*) at the N-terminus and the C2H2 Zn-fingers. (**B**) Domain architecture of Scm3 in *C. albicans* reveals a 623 aa long polypeptide consisting of the conserved Scm3 domain (aa 71-129) and three separate C2H2 zinc finger clusters (aa 407-570) (Aravind, Iyer et al. 2007).

**Table S1** (related to Figure 3, Figure S3). **Parameters used in simulation for polymer modelling of chromosomes in *C. albicans***

| **Simulation parameter** | **Values** |
|---|---|
| Number of beads, N | 1432 |
| Spring stiffness, $k_s$ | 100 $k_BT/l_0^2$ |
| Spring stiffness, $k_c$ | 500 $k_BT/l_0^2$ |
| Interaction strength, $E_{ij}$ | 1 $k_BT/l_0^2$ |
| Natural length $l_0$ | 1 $l_0$ |
| Radius of sphere, $R_s$ | $12l_0$ |

**Table S2** (related to Figure 4, Figure S4). **Chromosomal coordinates for Orc4 binding at centromeres based on *C. albicans* Assembly 21**

| **Chromosome** | **CENPA binding region Coordinates (length)** | **Orc4 binding region Coordinates (length)** |
|---|---|---|
| 1 | Ca21Chr1 15662315-1566930 (4616 bp) | Ca21Chr1 1562748- 1566244 (3497 bp) |
| 2 | Ca21Chr2 1925206- 1929688 (4483 bp) | Ca21Chr2 1926183- 1929443 (3261 bp) |
| 3 | Ca21Chr3 822762-827727 (4966 bp) | Ca21Chr3 823057- 826863 (3807 bp) |
| 4 | Ca21Chr4 991382-996030 (4649 bp) | Ca21Chr4 992010- 995522 (3513 bp) |
| 5 | Ca21Chr5 4673814-472497 (5114 bp) | Ca21Chr5 468552- 471618 (3067 bp) |
| 6 | Ca21Chr6 979686-984007 (4322 bp) | Ca21Chr6 980541- 983910 (3370 bp) |
| 7 | Ca21Chr7 425129- 431652 (6524 bp) | Ca21Chr7 425910- 429297 (3388 bp) |
| R | Ca21ChrR 1742833- 1748598 (5766 bp) | Ca21ChrR 1743951- 1747274 (3324 bp) |

**REAGENT TABLE**

| Reagent or resource | Source | Identifier |
|---|---|---|
| **Chemicals** | | |
| CaOrc4 peptide | GeneMed Synthesis, USA | Custom synthesized |
| Freund's complete adjuvant | Sigma | Cat. no. F5881 |
| Freund's incomplete adjuvant | Sigma | Cat. no. F5506 |
| Beta-mercaptoethanol | HiMedia | Cat. no. MB041 |
| Hydroxyurea | HiMedia | Cat no. 6487 |
| Dimethyl sulfoxide (DMSO) | Sigma | Cat. no. D2650 |
| Nocodazole | Sigma | CAS 31430-18-9 |
| Doxycycline hyclate | Sigma | Cat. no. D9891 |
| Super Signal West Pico chemiluminescent substrate | Thermo Scientific | Cat. no. 34080 |
| DAPI | Sigma | Cat. no. 10236276001 |
| Lysing enzyme | Sigma | Cat. no. L1412 |
| Protein A-Sepharose beads | Sigma | Cat. no. P3391 |
| Nourseothricin | Werner Bioagents | CAS 96736-11-7 |
| SensiFast SYBR kit | Bioline | Cat. no. BIO-98020 |
| **Antibodies** | | |
| Polyclonal anti-Protein A | Sigma | Cat. no. P2921 |
| Monoclonal anti-PSTAIRE | Abcam | Cat. no. 9866 |
| Monoclonal anti-GFP | Roche | Cat. no. 11814460001 |
| Goat anti-rabbit IgG-HRP | Bangalore Genei | Cat. no. 105499 |
| Goat anti-mouse IgG-HRP | Bangalore Genei | Cat. no. HP06 |
| AlexaFluor goat anti-rabbit IgG 568 | Invitrogen | Cat. no. 11011 |
| AlexaFluor goat anti-rabbit IgG 488 | ThermoFischer | Cat no. A27034 |
| **Experimental models (*C. albicans* strains used)** | | |
| **Name (Description)** | **Genotype** | **Reference** |
| SC5314 | Wild type | (Aszalos, Robison et al. 1968) |
| YJB8675 | *Δura3::imm434/Δura3::imm434,Δhis1::hisG/ Δhis1::hisG ,Δarg4::hisG/Δarg4::hisG, CSE4-GFP-CSE4/CSE4* | (Joglekar, Bouck et al. 2008) |
| CaLS301 (*MCM2* heterozygous null (SN148) | *Δura3::imm434/Δura3::imm434,Δhis1::hisG/ Δhis1::hisG ,Δarg4::hisG/Δarg4::hisG,* | This study |

| | | |
|---|---|---|
| | *Δleu2::hisG/Δleu2::hisG, MCM2::NAT/MCM2* | |
| CaLS302 (*MCM2* heterozygous null (SN148)) | *Δura3::imm434/Δura3::imm434,Δhis1::hisG/ Δhis1::hisG ,Δarg4::hisG/Δarg4::hisG, Δleu2::hisG/Δleu2::hisG, MCM2::FRT/MCM2* | This study |
| CaLS303 (*mcm2* conditional mutant (SN148)) | *Δura3::imm434/Δura3::imm434,Δhis1::hisG/ Δhis1::hisG,Δarg4::hisG/Δarg4::hisG,Δleu2:: hisG/Δleu2::hisG, MCM2:: FRT/MET3prMCM2* | This study |
| CaLS304 (*mcm2* conditional mutant (SN148)) | *Δura3::imm434/Δura3::imm434,Δhis1::hisG/ Δhis1::hisG ,Δarg4::hisG/Δarg4::hisG, Δleu2::hisG/Δleu2::hisG, MCM2:: FRT/MET3prMCM2* | This study |
| CaLS305 (*mcm2* conditional mutant (SN148)) | *Δura3::imm434/Δura3::imm434,Δhis1::hisG/ Δhis1::hisG ,Δarg4::hisG/Δarg4::hisG, Δleu2::hisG/Δleu2::hisG, MCM2:: FRT/MET3prMCM2* | This study |
| CaLS306 (*mcm2* conditional mutant (SN148) CENPA-Prot A) | *Δura3::imm434/Δura3::imm434,Δhis1::hisG/ Δhis1::hisG ,Δarg4::hisG/Δarg4::hisG, Δleu2::hisG/Δleu2::hisG, MCM2:: FRT/MET3prMCM2 CSE4 TAP(HIS)/CSE4* | This study |
| CaLS307 (*mcm2* conditional mutant (SN148) CENPA-Prot A) | *Δura3::imm434/Δura3::imm434,Δhis1::hisG/ Δhis1::hisG ,Δarg4::hisG/Δarg4::hisG, Δleu2::hisG/Δleu2::hisG, MCM2:: FRT/MET3prMCM2 CSE4 TAP(HIS)/CSE4* | This study |
| CaLS308 (*mcm2* conditional mutant (SN148) CENPA-Prot A) | *Δura3::imm434/Δura3::imm434,Δhis1::hisG/ Δhis1::hisG ,Δarg4::hisG/Δarg4::hisG, Δleu2::hisG/Δleu2::hisG, MCM2:: FRT/MET3prMCM2 CSE4 TAP(HIS)/CSE4* | This study |
| CaLS309 (*MCM2* heterozygous null (8675)) | *Δura3::imm434/Δura3::imm434,Δhis1::hisG/ Δhis1::hisG ,Δarg4::hisG/Δarg4::hisG, MCM2::NAT/MCM2 CSE4-GFP-CSE4/CSE4* | This study |
| CaLS310 (*MCM2* heterozygous null (8675)) | *Δura3::imm434/Δura3::imm434,Δhis1::hisG/ Δhis1::hisG ,Δarg4::hisG/Δarg4::hisG, MCM2::NAT/MCM2 CSE4-GFP-CSE4/CSE4* | This study |
| CaLS311 (*mcm2* conditional mutant (8675)) | *Δura3::imm434/Δura3::imm434,Δhis1::hisG/ Δhis1::hisG ,Δarg4::hisG/Δarg4::hisG, MCM2::FRT/MET3prMCM2 CSE4-GFP CSE4/CSE4* | This study |
| CaLS312 (*mcm2* conditional mutant (8675)) | *Δura3::imm434/Δura3::imm434,Δhis1::hisG/ Δhis1::hisG ,Δarg4::hisG/Δarg4::hisG, MCM2::FRT/MET3prMCM2 CSE4-GFP-CSE4/CSE4* | This study |
| CaLS313 (*mcm2* conditional mutant (8675)) | *Δura3::imm434/Δura3::imm434,Δhis1::hisG/ Δhis1::hisG ,Δarg4::hisG/Δarg4::hisG,* | This study |

| | | |
|---|---|---|
| | *MCM2::FRT/MET3prMCM2 CSE4-GFP-CSE4/CSE4* | |
| CaLS320 (*ORC4* heterozygous null (SN148)) | *Δura3::imm434/Δura3::imm434,Δhis1::hisG/Δhis1::hisG ,Δarg4::hisG/Δarg4::hisG, Δleu2::hisG/Δleu2::hisG, ORC4::NAT/ORC4* | This study |
| CaLS321 (*ORC4* heterozygous null (SN148)) | *Δura3::imm434/Δura3::imm434,Δhis1::hisG/Δhis1::hisG ,Δarg4::hisG/Δarg4::hisG, Δleu2::hisG/Δleu2::hisG, ORC4::NAT/ORC4* | This study |
| CaLS322 (*orc4* conditional mutant (SN148)) | *Δura3::imm434/Δura3::imm434,Δhis1::hisG/Δhis1::hisG ,Δarg4::hisG/Δarg4::hisG, Δleu2::hisG/Δleu2::hisG, ORC4::FRT/MET3prORC4* | This study |
| CaLS323 (*orc4* conditional mutant (SN148)) | *Δura3::imm434/Δura3::imm434,Δhis1::hisG/Δhis1::hisG ,Δarg4::hisG/Δarg4::hisG, Δleu2::hisG/Δleu2::hisG, ORC4::FRT/MET3prORC4* | This study |
| CaLS324 (*orc4* conditional mutant (SN148)) | *Δura3::imm434/Δura3::imm434,Δhis1::hisG/Δhis1::hisG ,Δarg4::hisG/Δarg4::hisG, Δleu2::hisG/Δleu2::hisG, ORC4::FRT/MET3prORC4* | This study |
| CaLS325 (*orc4* conditional mutant (SN148) CENPA-Prot A) | *Δura3::imm434/Δura3::imm434,Δhis1::hisG/Δhis1::hisG ,Δarg4::hisG/Δarg4::hisG, Δleu2::hisG/Δleu2::hisG, ORC4:: FRT/MET3prORC4 CSE4 TAP(HIS)/CSE4* | This study |
| CaLS326 (*orc4* conditional mutant (SN148) CENPA-Prot A) | *Δura3::imm434/Δura3::imm434,Δhis1::hisG/Δhis1::hisG ,Δarg4::hisG/Δarg4::hisG, Δleu2::hisG/Δleu2::hisG, ORC4:: FRT/MET3prORC4 CSE4 TAP(HIS)/CSE4* | This study |
| CaLS327 (*orc4* conditional mutant (SN148) CENPA-Prot A) | *Δura3::imm434/Δura3::imm434,Δhis1::hisG/Δhis1::hisG ,Δarg4::hisG/Δarg4::hisG, Δleu2::hisG/Δleu2::hisG, ORC4:: FRT/MET3prORC4 CSE4 TAP(HIS)/CSE4* | This study |
| CaLS328 (*ORC4* heterozygous null (8675) | *Δura3::imm434/Δura3::imm434,Δhis1::hisG/Δhis1::hisG ,Δarg4::hisG/Δarg4::hisG, ORC4:NAT/ORC4 CSE4-GFP-CSE4/CSE4* | This study |
| CaLS329 (*ORC4* heterozygous null (8675) | *Δura3::imm434/Δura3::imm434,Δhis1::hisG/Δhis1::hisG ,Δarg4::hisG/Δarg4::hisG, ORC4:NAT/ORC4 CSE4-GFP-CSE4/CSE4* | This study |
| CaLS330 (*orc4* conditional mutant (8675)) | *Δura3::imm434/Δura3::imm434,Δhis1::hisG/Δhis1::hisG ,Δarg4::hisG/Δarg4::hisG, ORC4::FRT/MET3prORC4 CSE4-GFP-CSE4/CSE4* | This study |
| CaLS331 (*orc4* conditional mutant (8675)) | *Δura3::imm434/Δura3::imm434,Δhis1::hisG/Δhis1::hisG ,Δarg4::hisG/Δarg4::hisG, ORC4::FRT/MET3prORC4 CSE4-GFP-CSE4/CSE4* | This study |

| | | |
|---|---|---|
| CaLS332 (*orc4* conditional mutant (8675)) | *Δura3::imm434/Δura3::imm434,Δhis1::hisG/ Δhis1::hisG ,Δarg4::hisG/Δarg4::hisG, ORC4::FRT/MET3prORC4 CSE4-GFP-CSE4/CSE4* | This study |
| CAKS3b (CENPA under *PCK1* promoter) | *Δura3::imm434/ Δura3::imm434 Δhis1::hisG/ Δhis1::hisG Δarg4::hisG/ Δarg4::hisG CSE4::PCK1prCSE4/ cse4::hisG:URA:hisG* | (Sanyal and Carbon 2002) |
| LSK446 (CEN7 deletion) | *Δura3::imm434/Δura3::imm434, Δhis1::hisG/Δhis1::hisG ,Δarg4::hisG/Δarg4::hisG, CSE4/CSE4-GFP-CSE4 CEN7/CEN7::HIS1* | (Sreekumar, Jaitly et al. 2019) |
| SBC189 (*cdc15* mutant) | *ura3Δ::imm434/ura3Δ::imm434 ade2Δ::hisG/ade2Δ::hisG ENO1/eno1::ENO1-tetR-ScHAP4AD-3xHA-ADE2 URA3-TETp-CDC15/cdc15Δ::dpl200* | (Bates 2018) |
| CaKS107 (Mcm2-TAP) | *Δura3::imm434/Δura3::imm434,Δhis1::hisG/ Δhis1::hisG ,Δarg4::hisG/Δarg4::hisG, MCM2-TAP(NAT)/MCM2* | This study |
| CaLS334 (Mcm2-TAP) | *Δura3::imm434/Δura3::imm434,Δhis1::hisG/ Δhis1::hisG ,Δarg4::hisG/Δarg4::hisG, MCM2-TAP(NAT)/ MCM2::URA3* | This study |
| CaLS335 | *Δura3::imm434/Δura3::imm434,Δhis1::hisG/ Δhis1::hisG ,Δarg4::hisG/Δarg4::hisG, MCM2-TAP(NAT)/ MCM2::FRT* | This study |
| CaLS336 | *Δura3::imm434/Δura3::imm434,Δhis1::hisG/ Δhis1::hisG ,Δarg4::hisG/Δarg4::hisG, MCM2-TAP(NAT)/ MCM2::FRT* | This study |
| CaLS337 | *Δura3::imm434/Δura3::imm434,Δhis1::hisG/ Δhis1::hisG ,Δarg4::hisG/Δarg4::hisG, MCM2-TAP(NAT)/ MCM2::FRT* | This study |
| CaLS338 (MET3prMcm2-TAP/mcm2) | *Δura3::imm434/Δura3::imm434,Δhis1::hisG/ Δhis1::hisG ,Δarg4::hisG/Δarg4::hisG, MET3pr(URA3)MCM2-TAP(NAT)/ MCM2::FRT* | This study |
| CaLS339 (MET3prMcm2-TAP/mcm2) | *Δura3::imm434/Δura3::imm434,Δhis1::hisG/ Δhis1::hisG ,Δarg4::hisG/Δarg4::hisG, MET3pr(URA3)MCM2-TAP(NAT)/ MCM2::FRT* | This study |
| CaLS340 (MET3prMcm2-TAP/mcm2) | *Δura3::imm434/Δura3::imm434,Δhis1::hisG/ Δhis1::hisG ,Δarg4::hisG/Δarg4::hisG, MET3pr(URA3)MCM2-TAP(NAT)/ MCM2::FRT* | This study |
| CaAB1(Scm3 heterozygous) | Δ*ura3*::imm434/Δ*ura3*::imm434 Δ*his1*::hisG/ Δ*his1*::hisG Δ*arg4*::hisG/ Δ*arg4*::hisG Δleu2::hisG/ Δleu2::hisG *SCM3/scm3:NAT-Flp* | This study |

| | | |
|---|---|---|
| CaAB2 (Scm3 heterozygous) | Δ*ura3::*imm434/Δ*ura3::*imm434 Δ*his1::*hisG/ Δ*his1::*hisG Δ*arg4::*hisG/ Δ*arg4::*hisG Δleu2*::*hisG/ Δleu2*::*hisG *SCM3/scm3:FRT* | This study |
| CaAB3 (Met3prScm3) | Δ*ura3::*imm434/Δ*ura3::*imm434 Δ*his1::*hisG/ Δ*his1::*hisG Δ*arg4::*hisG/ Δ*arg4::*hisG Δleu2*::*hisG/ Δleu2*::*hisG *MET3p-SCM3-URA3/scm3:FRT* | This study |
| CaAB4 (Met3prScm3) | Δ*ura3::*imm434/Δ*ura3::*imm434 Δ*his1::*hisG/ Δ*his1::*hisG Δ*arg4::*hisG/ Δ*arg4::*hisG Δleu2*::*hisG/ Δleu2*::*hisG *MET3p-SCM3-URA3/scm3:FRT* | This study |
| CaAB5 (Scm3 heterozygous) | *CSE4/CSE4-GFP-CSE4 scm3::NAT-Flp/SCM3* | This study |
| CaAB6 (Scm3 heterozygous) | *CSE4/CSE4-GFP-CSE4 scm3::NAT-FRT/SCM3* | This study |
| CaAB7 (Met3prScm3) | *CSE4/CSE4-GFP-CSE4 MET3p-SCM3-URA3/scm3:FRT* | This study |
| CaKS102 (Cse4-ProtA) | *Δura3::imm434/Δura3::imm434,Δhis1::hisG/ Δhis1::hisG, Δarg4::hisG/Δarg4::hisG, Δleu2::hisG/Δleu2::hisG CENP-A/CENP-A-TAP(URA3)* | (Mitra, Gomez-Raja et al. 2014) |
| CaAB8 (Scm3 heterozygous) | *CENP-A/CENP-A-TAP(URA3) scm3::NAT-Flp/SCM3* | This study |
| CaAB9 (Scm3 heterozygous) | *CENP-A/CENP-A-TAP(URA3) scm3::FRT/SCM3* | This study |
| CaNV52 (Met3Scm3 in Cse4-ProtA) | *CENP-A/CENP-A-TAP(URA3) MET3p-SCM3-URA3/scm3:FRT* | This study |
| CaNV50 (Scm3-2xGFP) | *Δura3::imm434/Δura3::imm434 Δhis1::hisG/ Δhis1::hisG Δarg4::hisG/ Δarg4::hisG Δleu2::hisG/ Δleu2::hisG SCM3-2xGFP-URA3/SCM3* | This study |
| CaNV51 (Scm3-2xGFP/Ndc80-RFP) | *Δura3::imm434/Δura3::imm434 Δhis1::hisG/ Δhis1::hisG Δarg4::hisG/ Δarg4::hisG Δleu2::hisG/ Δleu2::hisG SCM3-2xGFP-URA3/SCM3 NDC80-RFP/NDC80* | This study |
| **Recombinant DNA** | | |
| **Plasmid name** | **Description** | **Reference** |
| pSFS2a | Recyclable SAT-flipper cassette | (Reuss, Vik et al. 2004) |
| pLSK1 | *ORC4* upstream sequence cloned in pSFS2a | This study |
| pLSK2 | Deletion cassette for ORC4 in pSFS2a | This study |
| pCaDIS | Plasmid for promoter replacement with MET3pr | (Care, Trevethick et al. 1999) |
| pLSK3 | *ORC4* N-terminus cloned in pCaDIS | This study |
| pLSK4 | *MCM2* upstream sequence cloned in pSFS2a | This study |
| pLSK5 | Deletion cassette for *MCM2* in pSFS2a | This study |

| | | |
|---|---|---|
| pLSK6 | Deletion cassette for *MCM2* using recyclable *URA3* | |
| pLSK7 | *MCM2* N-terminus cloned in pCaDIS | This study |
| pASB1 | *SCM3* upstream sequence cloned in pSFS2a | This study |
| pASB2 | Deletion cassette for *SCM3* in pSFS2a | This study |
| pASB3 | N-terminus of Scm3 cloned in pCaDIS | This study |
| pGFP-URA3 | GFP orf cloned in pBS-URA3 | (Chatterjee, Sankaranarayanan et al. 2016) |
| pNV31 | GFP tagging plasmid for Scm3 | This study |
| pNV32 | 2xGFP tagging plasmid for Scm3 | This study |
| pNdc80-RFP-ARG4 | RFP tagging plasmid for Ndc80 | (Varshney and Sanyal 2019) |
| **Oligonucleotides** | | |
| **Name** | **Sequence** | **Description** |
| ORC4_13 | CGGGGTACCTTGGTTTGTAAAAATGTTGTTTC | Deletion cassette for *ORC4* |
| ORC4_14 | CCGCTCGAGAAATAGTTTTACTCTTGAGTTAGC | |
| ORC4_15 | TCCCCGCGGGTTATAGGTTGCTTTTAGTGC | |
| ORC4_16 | TCCCCGCGGGTTATAGGTTGCTTTTAGTGC | |
| ORC4_11 | CGCGGATCCATGAATTCACAGGACC | N term of Orc4 (For *MET3pr* cloning) |
| ORC4_12 | AACTGCAGTGCCATTTAACTCTTTTAAGGCG | |
| MCM2_13 | CGGGGTACCCTAATCCCATTTTGTTATGAATAT | Deletion cassette for *MCM2* |
| MCM2_14 | CCGCTCGAGGGTTGATTAAATAGTAATGTAATTAATAAAG | |
| MCM2_15 | TCCCCGCGGGTGATTAGTGGGTTATGG | |
| MCM2_16 | CGGAGCTCTGCATTCCAGATTATTTTCTG | |
| MCM2_11 | CGCGGATCCATGTCAAGTCCACCAGCTG | N term of Mcm2 (For *MET3pr* cloning) |
| MCM2_12 | AACTGCAGGCGTCTTCATCTTCATCATCGTC | |
| M1 | AATTTTATCCAGATTTGATATTATG | TAP tagging of Mcm2 by overlap PCR |
| M2 | TTCCATCTTCTCTTTTCCATCAAAGTATATTTCATAAATTTACTTC | |
| M3 | GAAGTAAATTTATGAAATATACTTTGATGGAAAAGAGAAGATGGAA | |
| M4 | TACAAACAATAACAATAACTATAACGATATCAAGCTTGCCTCGTC | |

| | | |
|---|---|---|
| M5 | GACGAGGCAAGCTTGATATCGTTATAG TTATTGTTATTGTTTGTA | |
| M6 | TTCAAGATATTATAAAATAGTCGAA | |
| CEN1 CORE RT1 | CAATCTAGCATTTCCTTCACACA | qPCR for CEN1 |
| CEN1 CORE RT2 | TGACGCAATGAAGTAGGTGAT | |
| CEN2 FP RT | CTCATTCGGAAGATTATAGTACTTGG | qPCR for CEN2 |
| CEN2 RP RT | CATAGTCAATACAATACGTCTTCTG | |
| CEN3 FP RT | CCTGTGTTGTAAATCAGATCAG | qPCR for CEN3 |
| CEN3 RP RT | CATCCCTTGCTCTATCTTATTCAC | |
| CEN4 FP RT | GCATTAACGTTCTGCTGTTCTTAG | qPCR for CEN4 |
| CEN4 RP RT | CTCACCGGAACAGACTGAAC | |
| CACH5F1 | CCCGCAAATAAGCAAACACT | qPCR for CEN5 |
| CACH5R1 | TTCATGGAAGAGGGGTTTCA | |
| CEN6 FP RT | CGATTGATCCATCACGATGG | qPCR for CEN6 |
| CEN6 RP RT | CTTTTAGTGAGGATGTATGGGATGC | |
| nCEN7-3 | GCATACCTGACACTGTCGTT | qPCR for CEN7 |
| nCEN7-4 | AACGGTGCTACGTTTTTTTA | |
| CENR FP RT | GGAGCCGCCTAAAACTTTTG | qPCR for CENR |
| CENR RP RT | CTATTGCCATCCAGGCTG | |
| 7S14 RTF | GGATGTTGAGTTCAAAGCCTG | qPCR of nCEN7-II (neocen at Chr7) |
| 7S14 RTR | CCAGCCAAATAATCTAGCTGC | |
| C1_FP | GGCATAGTGAATAATTCTATAAGCAC | qPCR analysis (Orc4 binding regions at chromosome arms) |
| C1_RP | CAAGCTATGTGATTTGACTTAATGAAC | |
| C2_FP | GGCAGCAATATTTGGTGC | |
| C2_RP | GCCAAGAGATACAAATCAAACAAC | |
| C3_FP | CTACATCGGACTTGTTGTGTC | |
| C3-RP | CATTCTGAAGGTTACAACATATGC | |
| C4_FP | CGGGGAATCGAACCCC | |
| C4-RP | CAAGTCAGCTAATTAGCTCAG | |
| C5_FP | GTGGCGATCACATCGG | |
| C5-RP | CATAACATTTGCAAGGCAAG | |
| C6_FP | CCTTCTTACGGTGTTGCTG | |
| C6_RP | GTGAGCAAAAGCGTTATATC | |
| C7-FP | CCCCCAGTCTTGTTTTTAC | |
| C7_RP | ACAATTTTTATACCCATTTTACTGC | |
| CR_RP | GAAAGCAATGACTTCATAACCTTG | |
| CR_RP | CAACAGAAACATGTCAAAGGG | |
| nLeu2-1 | GTACCGAAATTGTCAATGAAG | |
| nLeu2-2 | GTGGTGTTTGAAATCAAATTG | |
| Non-CEN7a | ACTCGCCTTCCCCTCCTTTAAATAG | ChIP PCR at non-CEN7 |
| Non-CEN7b | CCACTACTACGACTGTGGATTCACT | |
| ASB25 | AGTGGTACCCCAGACGACATCAGGTGT TTC | Deletion cassette for *SCM3* |
| ASB26 | CAGCTCGAGGCTATGATTTACGGCAAA CAC | |

| | | |
|---|---|---|
| ASB27 | TCCCCGCGGTTATGCGGTTTCTGGAGCAG | |
| ASB28 | ATCGAGCTCAATGGCTCAACAAATGATCTTG | |
| ASB34 | CGCGGATCCATGAATACCGGTATTGAAAATGAT | N -term cloning of SCM3 (for *MET3* promoter) |
| ASB35 | TGCACTGCAGAGATTGATAAGTTTCATCGTCG | |
| ASB36 | TTGTCACTTTCTTCCTGTTAAC | Confirmation of *MET3pr* integration |
| ASB37 | CTGCTCCAGAAACCGCATAA | |
| ASB38 | TCTATTATCCCTCGTGGTCAAG | confirmation of *scm3* deletion |
| NATmidF | TTAGAGACACAAACGAACAATGTACC | |
| NV158 | TCCCCGCGGACTTTCATCTCAAACTGAAGAA | Construction of *SCM3-GFP* cassette |
| NV159 | CGGACTAGTATTGAATAATTCATCTATTGATTCATA | |
| SR67 | CGCACTAGTATGAGTAAGGGAGAAGAACTTTTCAC | Construction of *SCM3-2xGFP* cassette |
| NV250 | CGCACTAGT TTTGTATAGTTCATCCATGCC | |

| **Software and algorithms used:** | |
|---|---|
| **Name** | **Source/ Reference** |
| Candida genome database | http://www.candidagenome.org/ |
| Integrative Genomics Viewer | http://software.broadinstitute.org/software/igv/ |
| ESPript 3.0 | http://espript.ibcp.fr/ESPript/ESPript/ |
| MACS2 | (Feng, Liu et al. 2012) |
| Bowtie | (Langmead, Trapnell et al. 2009, Langmead and Salzberg 2012) |
| DIVERSITY | (Mitra, Biswas et al. 2018) |
| HiCUP | (Wingett, Ewels et al. 2015) |
| DryHiC | (Vidal, le Dily et al. 2018) |
| LAMMPS | (Plimpton 1995) |
